# Huntingtin is an RNA-binding protein and participates in NEAT1-mediated paraspeckles

**DOI:** 10.1101/2024.02.07.579162

**Authors:** Manisha Yadav, Rachel J. Harding, Tiantian Li, Xin Xu, Terence Gall-Duncan, Mahreen Khan, Costanza Ferrari Bardile, Glen L. Sequiera, Shili Duan, Renu Chandrasekaran, Anni Pan, Jiachuan Bu, Tomohiro Yamazaki, Tetsuro Hirose, Panagiotis Prinos, Lynette Tippett, Clinton Turner, Maurice A. Curtis, Richard L.M. Faull, Mahmoud A. Pouladi, Christopher E. Pearson, Housheng Hansen He, Cheryl H. Arrowsmith

## Abstract

Huntingtin protein, mutated in Huntington disease, is implicated in nucleic acid-mediated processes, yet evidence for direct huntingtin-nucleic acid interaction is limited. Here we show wildtype and mutant huntingtin co-purify with nucleic acids, primarily RNA, and interact directly with G-rich RNAs in in vitro assays. Huntingtin RNA immunoprecipitation sequencing from patient-derived fibroblasts and neuronal progenitor cells expressing wildtype and mutant huntingtin revealed NEAT1 as a significantly enriched transcript. Altered NEAT1 levels were evident in Huntington’s disease cells and postmortem brain tissues, and huntingtin knockdown decreased NEAT1 levels. Huntingtin co-localized with NEAT1 in paraspeckles, and we identified a high-affinity RNA motif preferred by huntingtin. This study highlights NEAT1 as a novel huntingtin interactor, demonstrating huntingtin’s involvement in RNA-mediated functions and paraspeckle regulation.

**One-Sentence Summary:** HTT is an RNA-binding protein that interacts with G-rich sequences, including those in the paraspeckle lncRNA NEAT1.

## Introduction

Huntington’s disease (HD) is a rare autosomal dominant neurodegenerative disorder with a wide range of motor, cognitive, and psychological symptoms (*1*, *2*). HD is caused by expansions of a naturally occurring CAG (encoding polyglutamine) repeat tract in the Huntingtin (*HTT*) gene (*3*). The *HTT* gene has 5-35 CAG repeats in unaffected individuals while, mutant *HTT* (*mHTT*) contains ≥36 CAG repeats (*4*, *5*). CAG repeat length is the primary driver of age-of-onset in HD, with other genetic factors such as polymorphic variants within DNA repair genes and repeat tract purity, also influencing the age-of-onset and progression of the disease (*6*, *7*).

HTT is a 348 kDa HEAT-repeat protein thought to serve as a scaffold for protein-protein interactions (*8*, *9*). Among these, HAP40 is the only structurally and biophysically characterized protein interactor that forms a stable hetero-dimer with both HTT and mHTT (*10–12*). Notably, the N-terminal HEAT domain has a positively charged solvent-exposed surface, which has been postulated to act as a binding site for nucleic acids (*11*). Indirectly, HTT has been implicated by colocalization experiments in RNA transport (*13*, *14*) and may interact with its own mRNA (*15*).

Although evidence for direct interaction of HTT with RNA is limited, there is significant literature implicating HTT in gene expression and stress responses through mechanisms that could involve HTT-RNA interactions. Furthermore, RNA binding proteins like FUS consists of prion-like domains (PLDs) enriched in disordered regions with low sequence complexity, are also implicated in the formation of liquid-liquid phase separation (LLPS) (*16*). Similar to this, exon 1 of HTT (HTTex1), comprising polyQ and proline-rich prion-like domain (PLD) regions, is known to undergo LLPS and transition to form higher-ordered assemblies, both *in vitro* and in cells (*17*, *18*). Both wildtype (WT) and mutant HTTex1 associate with Ago2 in P-bodies, suggesting a potential role for HTT in RNA-mediated gene silencing (*19*). Additionally, HTT functions in the nucleus and subnuclear speckles, contributing to transcription repression and RNA processing (*20*). During oxidative stress, HTT translocates to the nucleus, colocalizing with SC35+ nuclear speckles (*21*). Associations with Caprin-1 and G3BP1, stress granule markers, could suggest a role for HTT in regulating RNA processing and translation under stress (*22*, *23*). These findings suggest HTT’s involvement in LLPS with proteins and RNA, potentially impacting cellular RNA-dependent processes such as gene regulation and stress response.

To better understand HTT’s potential role in RNA mediated processes we used biophysical, biochemical, and cell-based assays to investigate the interaction of HTT and mHTT with RNA. We demonstrate direct interaction between HTT and RNA, identifying NEAT1 as a major HTT-binding RNA in cells, supporting a role for HTT in RNA-mediated stress responses involving NEAT1-mediated nuclear paraspeckles.

## RESULTS

### HTT-HAP40 interacts with RNA *in vitro*

We previously reported biophysical and structural studies of recombinant HTT (polyQ length of 23, or Q23) and expanded mHTT (Q54), and their HTT-HAP40 complexes, a biologically important proteoform of HTT and an obligate interaction partner of HTT (*11*, *24*, *25*). An intriguing observation was that both HTT and mHTT samples, extracted from either insect or mammalian cells via one-step affinity purification, co-purified with large amounts of nucleic acid, and additional purification steps were required to yield highly pure (>98%) HTT protein **(fig. S1A)**.

Using a 5’-FAM labeled, single stranded (ss) RNA and 100 bp double stranded DNA (dsDNA) oligonucleotides with the same random sequence **(Table 1)**, we conducted electrophoretic mobility shift assays (EMSA) with fully purified, recombinant HTT-HAP40 proteins **(fig. S1A)**. We observed protein concentration-dependent band shifts indicating formation of a complex of HTT- HAP40 with ssRNA, while dsDNA of the same sequence produced no shift. **(Fig. 1A, and fig. S1B)**. Quantitative fluorescence polarization (FP) assays using the same oligonucleotides confirmed a consistent trend, with HTT-HAP40 binding to ssRNA (KD = 1 ± 0.6 µM), while no significant binding to dsDNA (KD > 15 µM, highest titration concentration) **(Fig. 1B, and fig. S1C)**. We further validated this finding using surface plasmon resonance (SPR) analysis, yielding a similar trend with KD values of 0.4 ± 0.2 µM and 2.0 ± 0.2 µM, for ssRNA and dsDNA, respectively **(Fig. 1C, and fig. S1D)**. Similar binding results were obtained for mHTT-HAP40 (Q54) compared to the wildtype protein for both ssRNA (KD = 0.5 ± 0.3 µM) and dsDNA (KD > 15 µM) oligonucleotide substrates. These findings suggest limited influence of the polyglutamine tract on the observed interactions.

**Fig 1.**
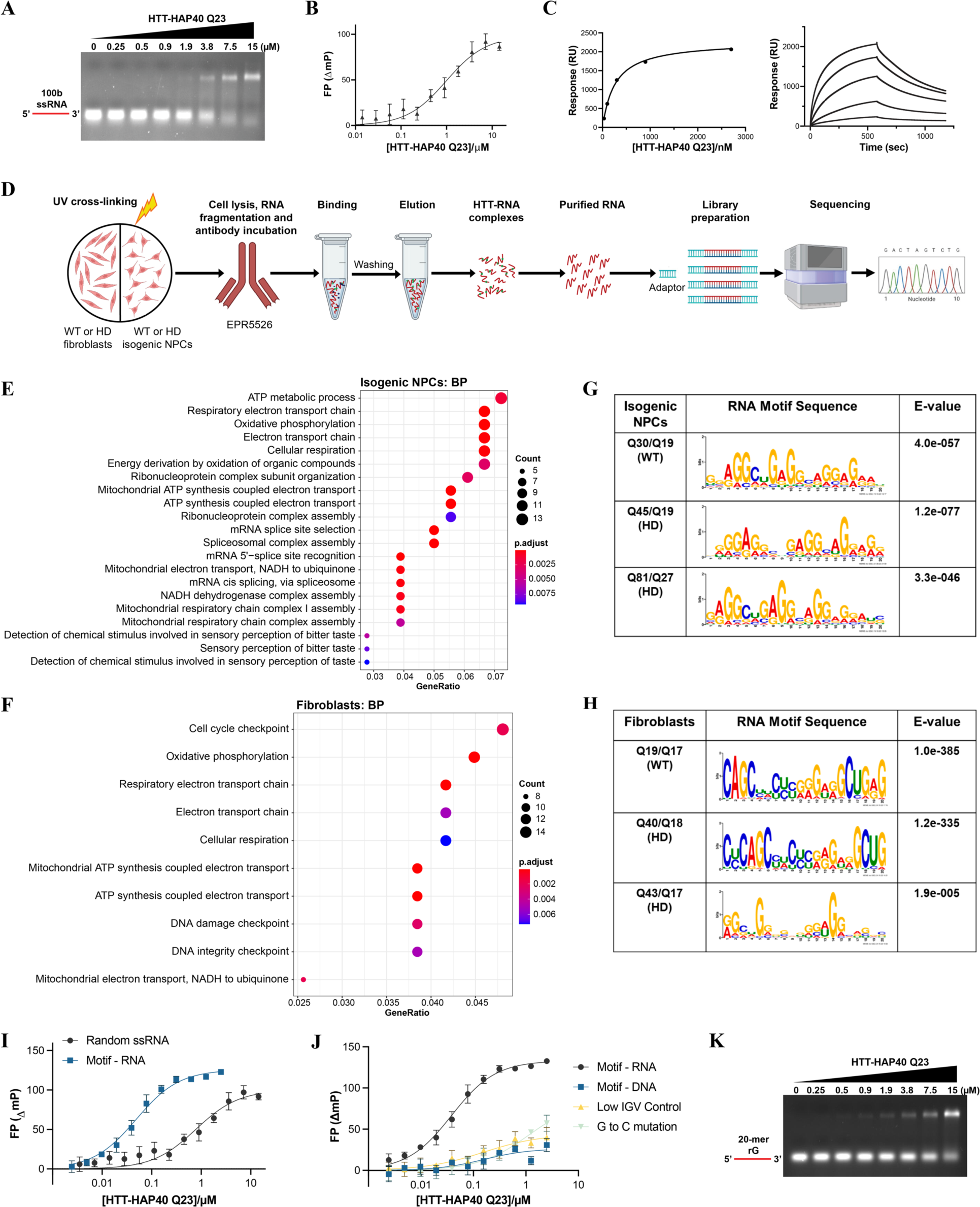
HTT protein binds RNA and prefers G-rich RNA sequences. (A) Representative EMSA images of increasing HTT-HAP40 Q23 protein (0–15 μM) binding with 1 µM of 100-mer random ssRNA. RNA is in red. EMSA, electrophoretic mobility shift assay. **(B)** Representative FP binding curve of HTT-HAP40 Q23 and 100-mer random ssRNA (KD = 1 ± 0.6 µM). **(C)** Representative SPR binding curve and sensorgram of HTT-HAP40 Q23 and 100-mer random ssRNA (KD = 0.4 ± 0.2 µM). **(D)** Schematic showing the protocol for HTT RIP-seq in isogenic NPCs and fibroblasts. **(E-F)** GO enrichment analysis for biological process (BP) in both WT and expanded isogenic NPCs (E) and fibroblasts (F). **(G-H)** MEME identified HTT-protein binding RNA motif sequences captured by isogenic NPC (**G**) and fibroblast (**H**) RIP-seq. **(I)** Representative FP binding curve of HTT-HAP40 Q23 and 100-mer random ssRNA (KD = 1 ± 0.6 µM) and motif RNA (KD = 40 ± 8.5 nM). **(J)** Representative FP binding curve of HTT-HAP40 Q23 and motif RNA, DNA form of motif RNA (KD > 15 µM), low IGV (KD > 15 µM) and G to C mutated RNA (KD > 15 µM). NC, not calculated, outside of range of protein concentrations tested. **(K)** Representative EMSA image of increasing HTT-HAP40 Q23 protein (0–15 μM) binding with 1 µM of 20-mer rG. RNA is in red. EMSA, electrophoretic mobility shift assay.

**Table 1.**
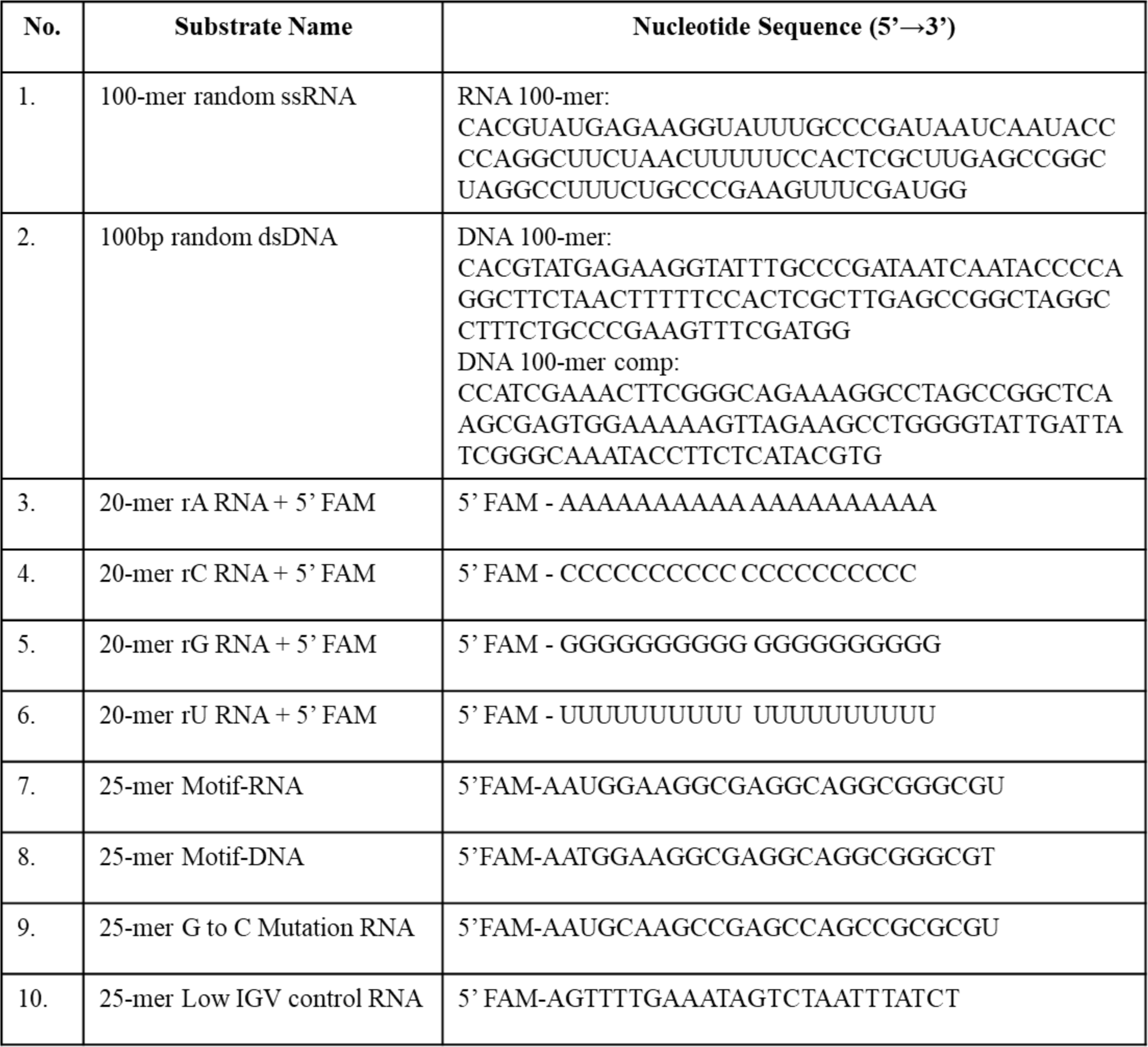
List of RNA and DNA substrates used in the study.

### HTT protein binds RNA and prefers G-rich RNA sequences

To gain insight into the endogenous RNAs bound by HTT in disease-relevant cells, we performed HTT RNA-immunoprecipitation sequencing (RIP-seq) in control and patient-derived fibroblast cells and isogenic neural progenitor cells (NPCs). Specifically, one WT (Q19/Q17) and two HD (Q40/Q18 and Q43/Q19) patient-derived fibroblast cell lines (*21*) and WT (Q30/Q19) and two isogenic HD NPC cell lines representing adult (Q45/Q19) and juvenile (Q81/Q27) disease-onset patients (*26*) were used. Cells were UV-crosslinked to form covalent bonds between RNA and protein, followed by immunoprecipitation (IP) of HTT and purification of its associated RNA transcripts and then sequencing of the prepared RNA libraries **(Fig. 1D, and fig. S2, A and B)**. Principle component analysis (PCA) showed grouping distinctions between input and IP samples, as well as the separation of WT and HD samples in both NPCs and fibroblasts **(fig. S2, C and D)**.

In total, 177 and 541 RNA transcripts were commonly and significantly enriched across all the WT and HD IP samples of NPCs **(fig. S3, A and B)** and fibroblasts **(fig. S3, C and D)**, respectively. A subset of 53 transcripts were common between the isogenic NPCs and fibroblasts, including NEAT1, ADARB1, EIF4A1, HOTAIRM1, BCL6, DOHH, CHKB as well as certain other RP11 transcripts, micro-RNAs, and mitochondrial-encoded RNAs. We found significant enrichment of HTT RNA transcript in WT NPCs, similar to reported earlier (*15*), however, it was not enriched in any other samples. Gene ontology (GO) enrichment analysis for biological process (BP) and cellular components (CC) showed that the enriched transcripts were associated with pathways involved in mitochondrial ATP (MT-ATP) synthesis-coupled electron transport, RNA splicing, spliceosome machinery, 5’-splice site recognition, and several other mitochondrial functional pathways such as oxidative-phosphorylation (OXPHOS), electron transport chain and cellular respiration **(Fig. 1, E and F, and fig. S3, E and F)**.

To delve deeper into HTT-RNA interactions, we sought to determine whether HTT exhibits a preference for binding to specific RNA sequence(s). MEME motif sequence analysis (*27*) identified a G-rich sequence HTT-binding RNA motif (GGAAGGCGAGGC) in both NPCs **(Fig. 1G)** and fibroblasts **(Fig. 1H)**. To evaluate direct binding of HTT to the identified G-rich RNA motif we used synthetic, 5’-FAM labeled, 25-mer oligonucleotides bearing the identified RNA motifs, in a fluorescence polarization (FP) assay where HTT proteins were titrated (**Table 1**). HTT- HAP40 Q23 exhibited binding to the identified RNA motif, (labeled as Motif-RNA) with a KD value of 40 ± 8.5 nM compared to the same 100b random ssRNA used above **(Fig. 1, A to C)** with a KD = 1 ± 0.6 µM **(Fig. 1I)**. We also assessed the binding affinity of HTT-HAP40 Q23 to the RNA motif sequence within a DNA backbone, (labeled as Motif-DNA) which indicated a much weaker binding affinity (KD= not calculated), supporting HTT’s preference for RNA over DNA for this motif **(Fig. 1J)**. We further tested a region of NEAT1 that had no peaks enriched in our RIP-seq data (labeled as low IGV Control), which also showed weak binding to HTT-HAP40 Q23 **(Fig. 1J).**

Considering the preferred RNA motifs were all G-rich, a sequence that is prone to secondary structures, we conducted circular dichroism (CD) spectroscopy analysis of the motifs **(Table 1)** to assess for potential secondary structures. We found that the motif with either an RNA or DNA backbone was likely formed into G-quadruplexes **(fig. S4A)**, which could indicate that HTT has a preference to bind to these structures. To test if HTT’s binding is influenced by G-quadruplex structured elements, we replaced three guanines (G’s) in the RNA motif with cytosines [(G**C**AAG**C**CGAG**C**C)] (labeled as G to C mutation); which would disrupt the ability of the sequence to form a G-quadruplex **(Table 1)**. This G to C mutation resulted in significantly reduced binding of HTT-HAP40 Q23 (KD = greater than highest concentration tested **(Fig. 1J)**.

To validate this finding, we conducted an EMSA assay using 20-mer ssRNA substrates (rG, rU, rC, and rA) **(Table 1)** with increasing concentrations of HTT-HAP40 Q23 protein. Our EMSA results revealed formation of RNA-protein complexes in the presence of rG **(Fig. 1K)**. However, no significant RNA-protein complex formation was observed in the presence of rU, rC, and rA substrates **(fig. S4, B to D)**. Thus, our study suggests that HTT-HAP40 Q23 protein exhibits a preference for binding G-rich RNA motifs, aligning with our motif analysis and FP assay results. These findings contribute to our understanding of the molecular interactions involving HTT- HAP40 Q23 and its RNAs, particularly those with G-quadruplex forming sequences.

### Long non-coding RNA NEAT1 is a highly enriched HTT-bound transcript

The long non-coding RNA (lncRNA) NEAT1 was consistently the topmost significantly enriched transcript across all IP samples from different genetic backgrounds **(Fig. 2, A to C, and fig. S5, A to C)**. Previous studies have highlighted the presence of abundant and conserved G-quadruplex motifs in NEAT1, with RNA-binding proteins such as NONO exhibiting specificity for these motifs (Simko et al., 2020). NEAT1 displayed a consistent enrichment pattern across all RIP-seq samples (**Fig. 2D**). We used RT-qPCR to quantify the enrichment of the NEAT1 transcript in the IP fractions compared to the cytoplasmic RNA encoding the 40S ribosomal protein RPS28, as well as U6, and MALAT1, which are exclusively nuclear transcripts. A robust signal was detected for NEAT1 transcripts in the IP fractions compared to the housekeeping genes RPS28, U6 and lncRNA MALAT1, used as a negative control **(Fig. 2E, and fig. S4, D and E)**.

**Fig 2.**
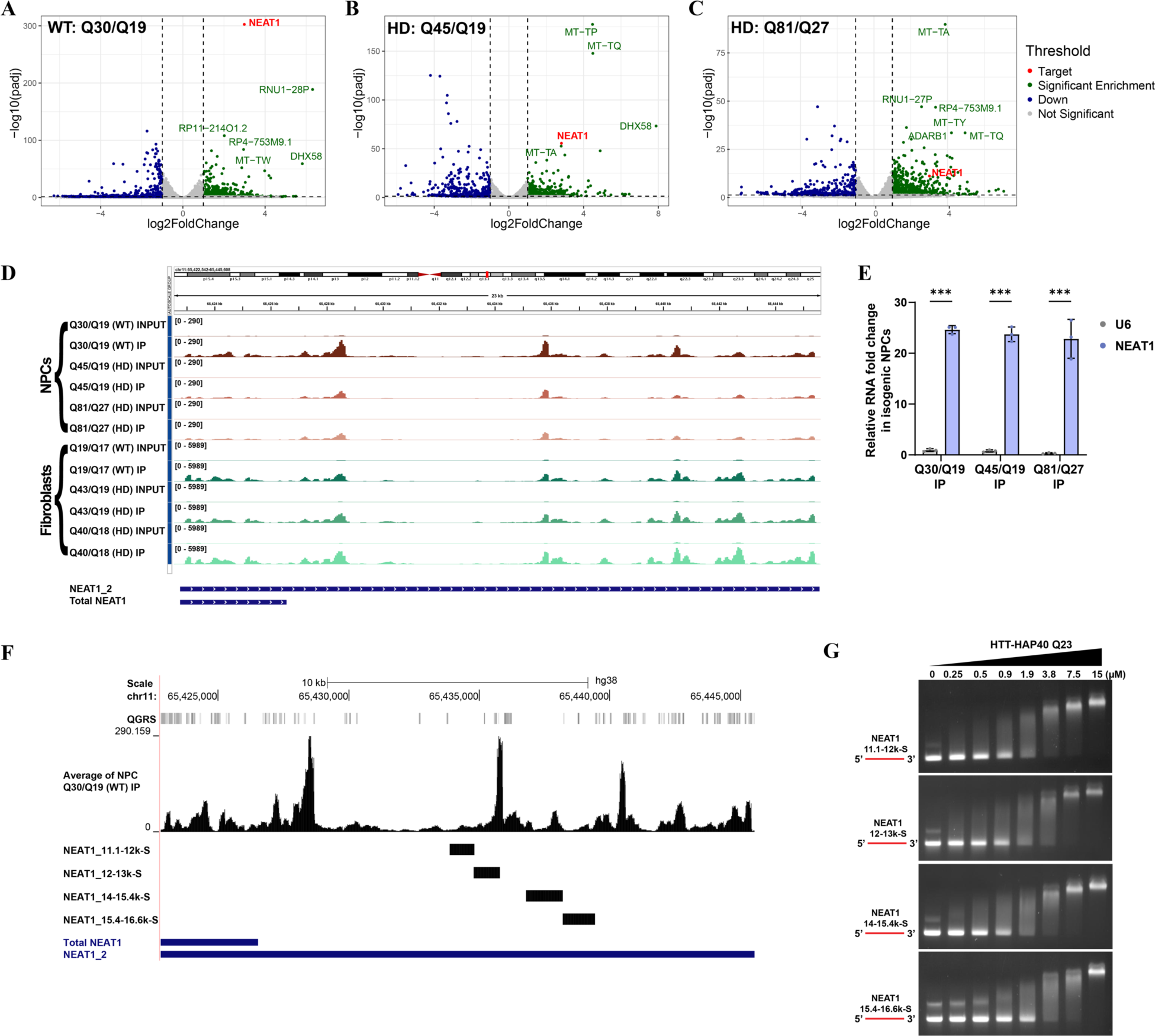
LncRNA NEAT1 as a novel HTT binding substrate. (A-C) Volcano plots showing lncRNA NEAT1 as a significantly enriched target in the WT and expanded isogenic NPCs IP samples (Log2 fold change cutoff: 1; p-value cutoff: 0.05). **(D)** Integrative Genomic Viewer snapshot showing the enrichment profile of NEAT1 across individual NPC and fibroblast IPs after autoscaling to their respective Inputs. **(E)** RT-qPCR validation of NEAT1 transcript enrichment in WT and expanded NPC IP samples compared to U6 control gene. Data was analyzed using 2-way ANOVA and shown as mean ± s.d.; *n* = 3. ***P < 0.001. **(F)** Isogenic NPCs Q30/Q19 IP data mapped to NEAT1. Quadruplex-forming G-rich sequences (QGRS) mapped to NEAT1 are shown as tall bars. *In vitro* transcribed NEAT1 fragments are mapped to the NEAT1 gene. **(G)** Representative EMSA images of increasing HTT-HAP40 Q23 protein (0–15 μM) binding with 250 nM of NEAT1 RNAs in red.

Given that the HTT-binding RIP-seq motif featured G-rich sequences **(Fig. 1, G and H**), and our recombinant HTT-HAP40 preferred G-rich ssRNA and G-quadruplexes (**Fig. 1, I to K**) we searched our NEAT1 RIP-seq data for for G-rich sequences. Using the QGRS mapper tool (https://bioinformatics.ramapo.edu/QGRS/index.php), employing parameters similar to Simko et al., we identified putative G-quadruplex-forming sequences in NEAT1_2. Analyzing NEAT1 enrichment profiles in NPC IP samples and QGRS data on the UCSC genome browser, we observed enrichment of HTT at multiple regions across NEAT1. Notably, these regions appeared to largely overlap with G-quadruplex-forming segments **(Fig. 2F)**. To further explore the interactions between HTT and NEAT1 RNA, we synthesized four *in vitro* transcribed sense RNA fragments of NEAT1 **(Fig. 2F)**, following the previously described protocol (*28*). Using an EMSA assay of the resultant transcripts, band shifts were observed in a concentration dependent manner with increasing concentration of HTT-HAP40 Q23 protein, providing further evidence of a direct interaction between HTT protein and lncRNA NEAT1 **(Fig. 2G)**.

### NEAT1 levels in NPCs, patient derived fibroblasts, and human post-mortem brain tissues are altered in HD

NEAT1 levels are reported to be dysregulated in various neurodegenerative diseases, including HD (*29–32*). Nevertheless, the association between NEAT1 and HD pathology remains unclear, with conflicting reports indicating either an increase or decrease in NEAT1 expression levels in various HD models (*32*, *33*).

The NEAT1 locus produces two isoforms: NEAT1_1 (short, ∼3.7kb) and NEAT1_2 (long, ∼22.7kb), **(Fig. 2, D and E)** with NEAT1_2 being an essential component of paraspeckles (*28*, *34*). We therefore assessed the levels of NEAT1 in the isogenic NPC allelic series, with RT-qPCR using NEAT1 isoform-specific primers. Because the NEAT1_1 sequence is identical to the 5’ sequence of the longer NEAT1_2, it is difficult to distinguish between the two (*35*). Therefore, primers that amplify the 5’ region of the gene will report on both NEAT1_1 and NEAT1_2 and such results are referred to as total NEAT1 throughout the text, whereas NEAT1_2 represents the long isoform (see also Fig. 2D, 2F). The qPCR results indicated that both total NEAT1 and NEAT1_2 levels were significantly lower in the polyQ-expanded NPCs compared to WT **(Fig. 3A)**. Similarly, qPCR of WT and HD fibroblast cell lines showed a significant reduction of NEAT1 in the HD cell lines compared to WT (Q21/Q18), with the homozygous HD cell line displaying significantly lower NEAT1 levels compared to the heterozygous HD cell lines **(Fig. 3B)**.

**Fig 3.**
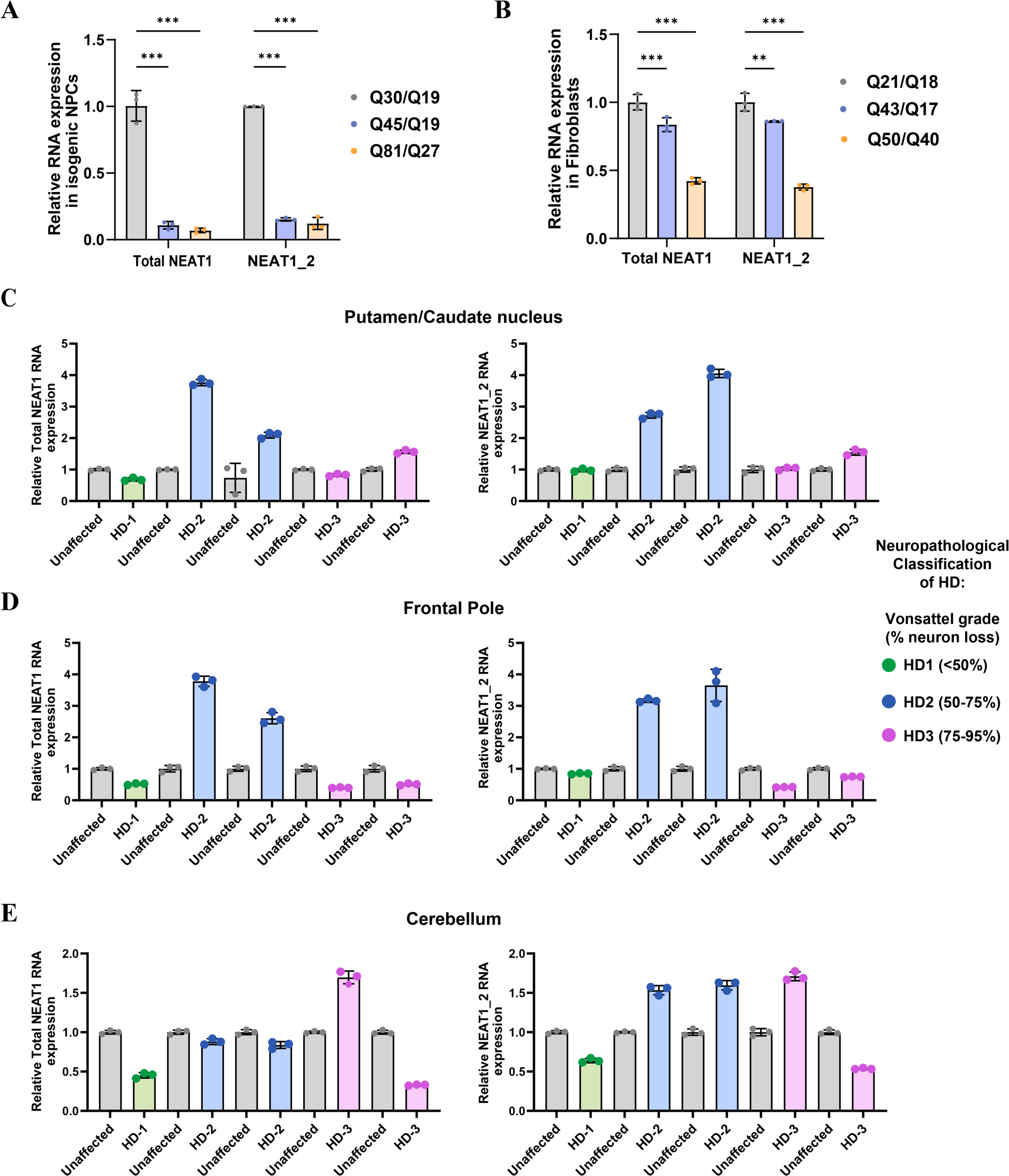
Altered levels of lncRNA NEAT1 across WT and expanded cell lines and HD human brain samples (A-B) RT-qPCR quantification of expression levels of total NEAT1 and long NEAT1 (NEAT1_2) isoform in WT and expanded isogenic NPCs (A) and patient-derived fibroblasts (B). U6 was used as control gene and data was analyzed using 2-way ANOVA. Data was analyzed using two-way ANOVA. Data are shown as mean ± s.d.; n = 3. ***P < 0.001, **P < 0.01. **(C-E)** RT-qPCR quantification of total NEAT1 and long NEAT1 (NEAT1_2) expression levels in putamen/caudate nucleus (C), frontal pole (D), and cerebellum (E) regions of human unaffected and HD patient (at different HD-grades) brain. Age and sex matched unaffected brain tissues for each region were used to normalize data (gray). *n* = 5 HD and unaffected individuals/group/tissue, 3 technical replicates/person. Dots indicate striatal neuropathological grade (HD 1–3). U6 was used as a control gene. See fig. S6 for an alternative figure of the same data. No statistics applied due to limited patient sample size.

Subsequently, we conducted qPCR using cDNA from three distinct brain regions (putamen/caudate nucleus, frontal pole, and cerebellum) derived from postmortem brain tissues of HD patients kindly provided from the Neurological Foundation Human Brain Bank in the Centre for Brain Research, University of Auckland with full consent from the donor families. These tissues were collected from HD patients diagnosed with different grades of disease **(Table 2)**. To normalize the RT-qPCR data, we utilized age- and sex-matched unaffected brain tissues **(Table 2)**. The qPCR results suggest a potential relationship between NEAT1 and human tissues from different HD grades, where we observed lower levels of both total NEAT1 and NEAT1_2 in HD grade 1, in all three brain regions. Interestingly, the NEAT1 levels seem to increase as the disease progresses from grade 1 to grade 2, especially the putamen and frontal pole **(Fig. 3, C to E, and fig. S6)**. This might potentially reflect the heightened stress on neurons. However, NEAT1 levels appear to subsequently decrease in grade 3, which could possibly be attributed to either neuronal loss or the presence of mutant HTT (mHTT) **(Fig. 3, C to E, and fig. S6)**. Larger sample sizes from different brain regions of unaffected and HD patients across different grades are required for robust statistical analysis and to draw any conclusions.

**Table 2.** Demographic characteristics of human postmortem brain tissue.

| Subject | Condition | Age<br>(years) | Sex<br>(M/F) | CAG | Region<br>sampled | COD | PMI<br>(hr) |
| --- | --- | --- | --- | --- | --- | --- | --- |
| **HD** |  |  |  |  |  |  |  |
| 1. | HD-1 | 32 | M | 17/47 | Put/CN,<br>CB, FP | Submandibular<br>squamous cell<br>carcinoma | 14 |
| 2. | HD-2 | 65 | M | 17/43 | Put/CN,<br>CB, FP | Renal failure | 14 |
| 3. | HD-2 | 70 | F | 17/42 | Put/CN,<br>CB, FP | HD | 8 |
| 4. | HD-3 | 64 | M | 27/42 | Put, CB,<br>FP | Pulmonary<br>thromboembolus | 18 |
| 5. | HD-3 | 63 | M | 22/43 | Put/CN,<br>CB, FP | Pulmonary<br>embolism | 16 |
| **Control** |  |  |  |  |  |  |  |
| 1. | Control | 22 | M | 11/23 |  | Asphyxia | 21 |
| 2. | Control | 60 | M | 10/17 | Put, CB,<br>FP | Ischaemic heart<br>disease | 17 |
| 3. | Control | 78 | F | 18/19 | Put/CN,<br>CB, FP | Aortic aneurysm | 20 |
| 4. | Control | 64 | M | 13/15 | Put, CB,<br>FP | Ischemic Heart<br>Disease | 15.5 |
| 5. | Control | 68 | M | 17/19 | Put, CB,<br>FP | Coronary<br>atherosclerosis | 22.5 |

*HD* Huntington's disease, *COD* cause of death, *PMI* postmortem interval, *Put* putamen, *CN* caudate nucleus, *CB* cerebellum, *FP* frontal pole

### HTT levels influence lncRNA NEAT1 levels

To understand whether there is a relationship between HTT and NEAT1 levels we knocked down HTT in WT (Q21/Q18) and two HD (Q43/Q17, Q57/Q17) fibroblast cell lines using both siRNAs **(fig. S7A, C to E)** and shRNAs **(Fig. 4, A to C).** HTT knockdown resulted in a reduction of both isoforms of NEAT1 in both WT and HD fibroblasts. We also assessed NEAT1 levels in two control cell lines; HEK293T cells expressing WT HTT and a derivative cell line in which full-length HTT was entirely knocked out (*36*) **(fig. S7, B to F)**, as well as an hTERT immortalized RPE1 cell line, which is Tet-inducible for HTT knockdown **(Fig. 3, A to D)**. Again, we observed that NEAT1 levels were reduced upon knockdown of HTT in RPE1 cells and in HEK293T HTT null cells compared to the parental cells, expressing normal HTT **(fig. S7F)**. Taken together these data suggest that the levels of HTT protein positively correlated with the abundance of NEAT1 lncRNA isoforms in a variety of cell lines.

**Fig 4.**
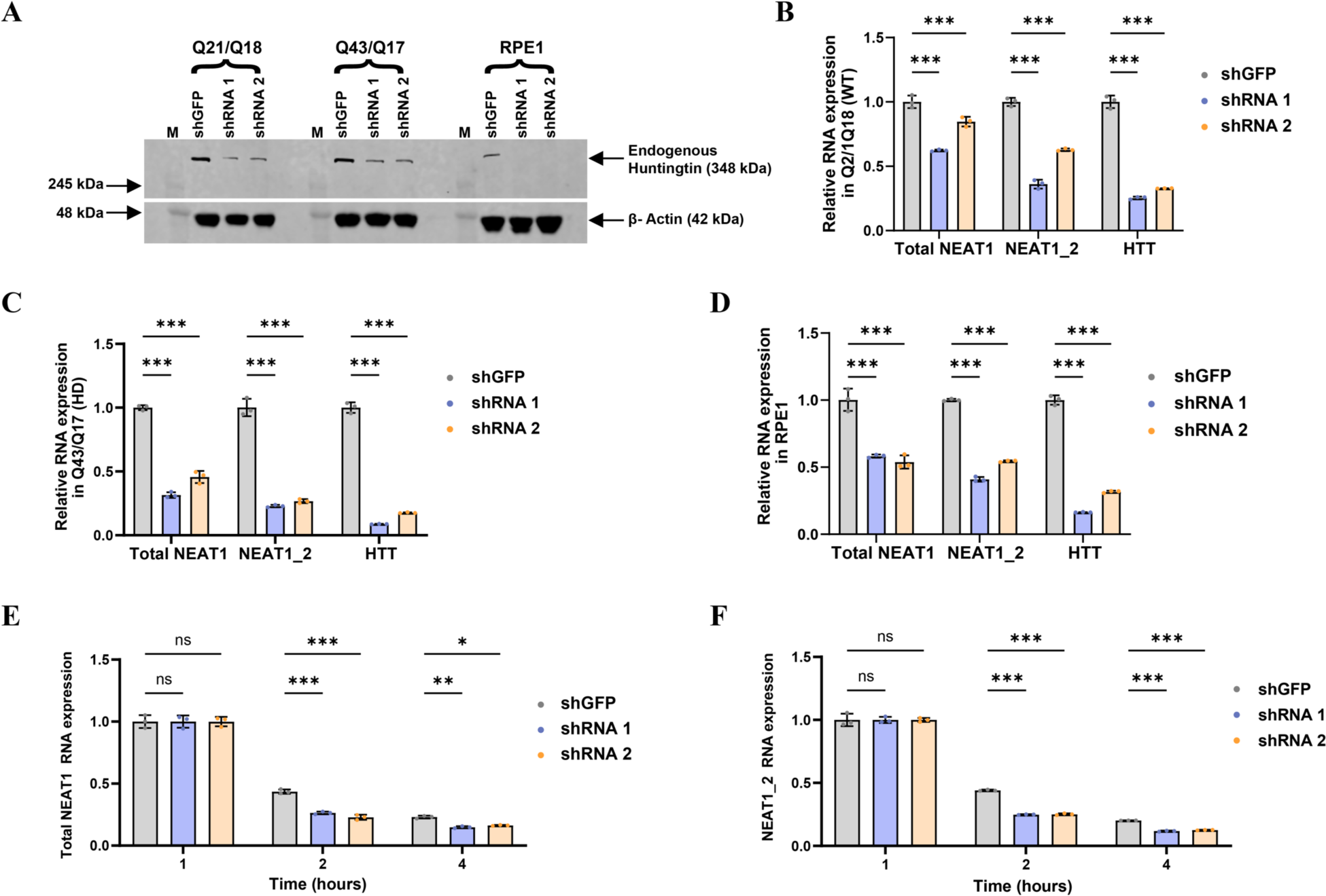
HTT knockdown reduces NEAT1 levels and HTT stabilizes lncRNA NEAT1. (A) Western blot analysis of HTT knockdown by shRNAs in WT (Q21/Q18), HD (Q43/Q17) fibroblast and RPE1 cell lines **(B-D)** RT-qPCR quantification of lncRNA NEAT1 isoforms upon HTT knockdown by shRNAs in WT (Q21/Q18) and HD (Q43/Q17) fibroblasts and RPE1 cell lines. **(E-F)** Graphs depicting the decay of Total NEAT1 **(E)** and NEAT1_2 **(F)** in shGFP control and shRNAs HTT-knockdown (KD) RPE1 cells following Actinomycin D treatment. U6 was used as control gene and data were analyzed using 2-way ANOVA. Data are shown as mean ± s.d.; *n* = 3. ***P < 0.001, **P < 0.01, *P < 0.05, ns, not significant.

We hypothesized that HTT protein may be contributing to the stability of NEAT1 lncRNA, thereby reducing its half-life under conditions of low HTT levels. To test this possibility, we used actinomycin-D to inhibit transcription in control and HTT knockdown RPE1 cells and assessed NEAT1 levels over time via qPCR. The abundance of both NEAT1 isoforms decreased dramatically within 2-4 hours with a significantly rapid reduction after HTT knockdown compared to the control cells expressing normal HTT **(Fig. 4, E and F)**. These findings suggest that HTT plays a crucial role in maintaining the stability of NEAT1 RNA following transcription.

### HTT and NEAT1 co-localize in RPE1 and fibroblast cell lines

To further understand the relationship between HTT and NEAT1, we used fluorescence *in situ* hybridization (FISH) and confocal microscopy to visualize and quantify NEAT1 in cells. As previously reported (Naganuma & Hirose, 2013) we observe both isoforms of NEAT1 are retained in the nucleus in discrete paraspeckles **(Fig. 5)**. Using RNA probes detecting either total NEAT1 (5’-NEAT1) or only NEAT1_2 (middle-NEAT1) we quantified the number of NEAT1 paraspeckles in WT fibroblast cells (Q21Q18) before and after HTT knockdown. The number of total NEAT1 and NEAT1_2 foci were significantly reduced after HTT protein knockdown in comparison to an shGFP control **(Fig. 5, A and B, and fig. S8, A and B)**. These data suggest that HTT contributes to NEAT1-mediated paraspeckle formation. Moreover, quantification of NEAT1 foci number and intensity using FISH revealed a significant reduction in the homozygous HD fibroblast cell line (Q50/Q40) compared to the WT (Q21/Q18) and heterozygous HD (Q43/Q17) fibroblasts **(Fig. 5C, and fig. S8, C to E)**, mirroring the qPCR results **(Fig. 3B),** and suggesting a potential deficit in NEAT1-mediated paraspeckles in homozygous HD cells.

**Fig 5.**
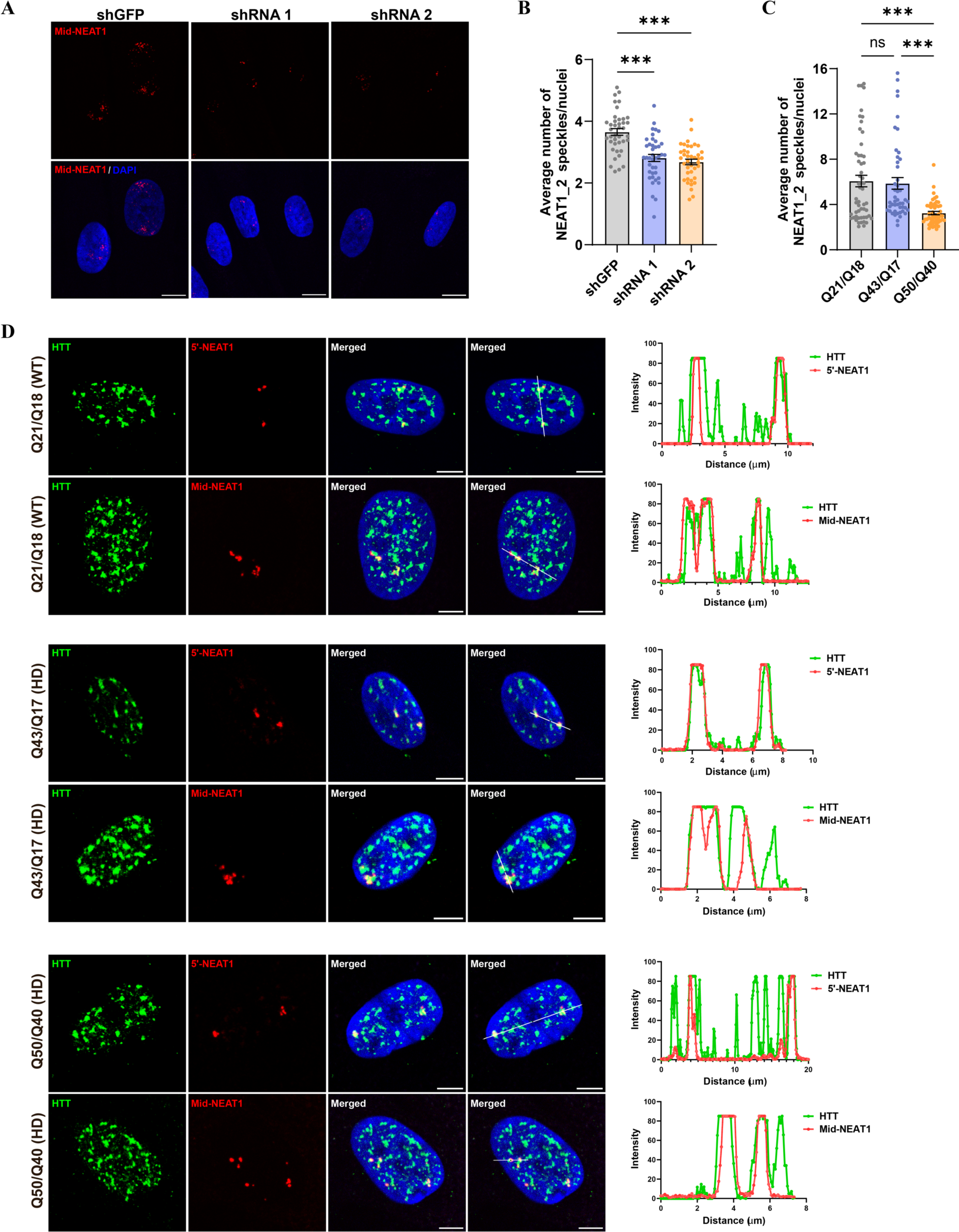
HTT co-localizes with NEAT1. (A-B) Representative images **(A)** and quantification **(B)** of NEAT1_2 positive foci in Q21Q18 fibroblast cell lines assessed by FISH. Scale bar = 10 μm. For statistical analysis of the NEAT1_2 RNA foci, CellProfiler software was used, and images were processed using ImageJ (Fiji app). **(C)** Quantification of NEAT1 levels in Q21Q18 (WT), Q43Q17 (heterozygous HD) and Q50Q40 (homozygous HD) fibroblast cell lines. Data represent mean ± sem. Data were analyzed by ordinary one-way ANOVA with Tukey’s test for multiple comparisons. ∗∗∗P < 0.001. **(D)** Phospho-N17 HTT antibody co-localizes with both 5’-NEAT1 and middle-NEAT1 probes in WT and HD patient derived fibroblast cells. Quantitation of HTT and NEAT1 co-localization performed using ImageJ (Fiji app). Scale bar indicates 5µm.

Our HTT-IP data suggests that HTT can bind to both isoforms of NEAT1 **(Fig. 2B)**. To confirm HTT-NEAT1 interaction, we combined immunofluorescence (IF) with RNA-FISH to examine whether HTT co-localizes with both isoforms, 5’- and middle-NEAT1 RNA within the cells. Nuclear HTT can also be visualized by immunofluorescence (IF) in distinct nuclear puncta, using an antibody that targets HTT phosphorylated serine residues 13 and 16 (phospho-N17) (*8*, *21*, *37*). We observed co-localization of HTT with both 5’- and middle-NEAT1 probes in WT (Q21/Q18), heterozygous HD (Q43/Q17) as well as homozygous HD (Q50/Q40) fibroblasts **(Fig. 5D, and fig. S9A).** Co-localization was also observed in RPE1 cells **(fig. S9B)**. These results further support direct HTT-NEAT1 interaction and suggest HTT participates in most NEAT1-positive nuclear PSs.

We performed Pearson’s correlation coefficient (PCC) analysis on co-localization images to assess differences between WT (Q21/Q18) and HD (Q43/Q17, Q50/Q40) fibroblast cell lines. The analysis revealed no significant distinctions between the WT and HD cell lines **(fig. S9C)**. Subsequently, we quantified the percentage of NEAT1 foci co-localized with phospho-N17 HTT in these cell lines, demonstrating that approximately 75-80% of NEAT1 foci co-localized with HTT foci. Both HD cell lines exhibited a trend toward reduced co-localization. While statistical significance was not consistently reached for the heterozygous HD (Q43/Q17) cells, the percentage co-localization in homozygous HD (Q50/Q40) cells was significantly decreased compared to WT **(fig. S9D)**. Again, a stronger phenotype in the HD (Q50/Q40) cells suggests a deficit in NEAT1 paraspeckle-related functions for mHTT.

A large proportion of HTT protein also formed nuclear foci without co-localizing with NEAT1. Since, HTT is also known to associate with SC35+ nuclear speckles (*21*), NEAT1 foci are reported to predominantly localize at the peripheries of SC35+ nuclear speckles (*38*, *39*). Therefore, we assessed if phospho-N17 HTT co-localized with both SC35+ nuclear speckles and NEAT1 paraspeckles. Confirming previous studies, we observed that phospho-N17 HTT co-localized with SC35+ nuclear speckles, in addition to NEAT1_2 foci in RPE1 cells in RPE1 cells. **(fig. S9E)**.

## Discussion

Our study demonstrates for the first time the ability of HTT protein to directly bind to RNA species, both *in vitro* and in cells, suggesting its potential involvement in RNA-mediated pathways. In our HTT RIP-seq experiments, we identified a G-rich RNA motif sequence with high affinity for HTT- HAP40, a major proteoform of HTT in cells (*25*). Our results also show that HTT interacts with NEAT1, which plays a crucial role as a structural scaffold for paraspeckles. Paraspeckles are dynamic, membrane-less nuclear bodies that influence various fundamental cellular functions and gene expression networks, particularly in response to stress induced by factors such as viral infections, proteasome inhibition, and mitochondrial stress (*16*, *29*, *40*). NEAT1 harbors conserved G-quadruplex motifs (*41*), aligning with our *in vitro* motif analysis and assays indicating the preference of HTT for G-rich sequences.

Our study has unveiled a reduction of NEAT1 levels in HD cell models compared to their WT counterparts. The reduction of NEAT1 levels in HD isogenic NPCs may be attributed to decreased expression of WT HTT protein, as previously observed in these cell lines with increasing polyQ repeats (*26*). This aligns with our finding that HTT knockdown reduces NEAT1 levels, underscoring an essential role of HTT in NEAT1-mediated paraspeckles. Furthermore, our results indicate a decrease in NEAT1 foci numbers, intensities, and percentage co-localization in HD fibroblasts, particularly in the homozygous HD cell line (Q50/Q40). Again, this decrease may be linked to the reduced levels of HTT protein in these cell lines, as previously reported (*21*). NEAT1 binding proteins such as NONO, SFPQ, and RBM14 are crucial for paraspeckle biogenesis and maintaining stability of the lncRNA, preventing it’s degradation (*29*). In our study, we discovered that HTT also plays a role in stabilizing both short and long NEAT1 isoforms, similarly aiding in preventing the degradation of NEAT1.

Recent research has highlighted the multifaceted functions of paraspeckles, including the regulation of various RNA-centric cellular processes like mRNA retention, A-to-I editing, mRNA cleavage, and protein sequestration (*35*, *40*, *42*, *43*). Our HTT RIP-seq analysis also revealed the enrichment of other transcripts such as ADARB1, EIF4A1 and DHX58 helicases, that are involved in stress response and other antiviral signaling pathways (*44–48*). Finally, we identified HTT RNA transcripts enriched in the WT NPCs (Q30), highlighting its association with its own mRNA as reported previously (*15*). Interestingly this enrichment was specific to the WT NPCs and not observed in other IP samples. Our analysis also unveiled the enrichment of various other lncRNAs such as HOTAIRM1, RP11, RP1, and RP13, along with mitochondrial encoded RNAs in all immunoprecipitated samples. These transcripts are linked to mitochondrial-functional pathways and RNA regulatory mechanisms, including splicing, transcription, and translation regulation. Cross-regulation between NEAT1-mediated paraspeckles and mitochondria has been reported earlier, where depletion of NEAT1 significantly impacts mitochondrial dynamics and function by altering the sequestration of mito-mRNAs within paraspeckles under stress conditions (*40*). Mitochondrial dysfunction is an early pathological mechanism in HD, where mHTT disrupts mitochondria releasing mito-RNAs into the cytoplasm. This, in turn, upregulates the innate immune response in the most vulnerable cell type, striatal spiny neurons (*49*, *50*). It is tempting to envision that mHTT may lead to compromised paraspeckle function, resulting in a reduced ability to cope with mitochondrial stress and potentially leading to cell death in relevant HD tissues. The function and mechanisms underlying these interactions, as well as potential differences in WT and HD conditions, warrants further investigation.

## Acknowledgments

We acknowledge family members who made this work possible through tissue donations and support. We acknowledge the Neurological Foundation Human Brain Bank, the Center for Brain Research, and the University of Auckland. We thank Drs. Marcy E. MacDonald and Ihn Sik Seong from Massachusetts General Hospital for a gift of HTT null HEK 293T cells. We thank Dr. Truant Ray for providing the TruHD fibroblast cell lines and N17-phospho HTT antibody.

## Author contributions

Conceptualization: MY, RJH, CHA, PP, HHH, CEP, TGD

Formal Analysis: MY, TL, XX

Investigation: MY, RJH, TL, XX, TGD, MK, CFB, GLS, SD, RC

Resources: CFB, GLS, TY, TH, LT, CT, MAC, RLMF, SD, RC, AP, JB

Funding acquisition: CHA, RJH, HHH, CEP, MAP

Supervision: CHA, RJH, HHH, CEP, MAP

Writing – original draft: MY, RJH, CHA, PP

Writing – review & editing: MY, RJH, CHA, HHH, CEP, MAP, PP, TGD, XX

## Data and materials availability

All data, code, and materials used in the analysis must be available in some form to any researcher for purposes of reproducing or extending the analysis. Include a note explaining any restrictions on materials, such as materials transfer agreements (MTAs). Note accession numbers to any data relating to the paper and deposited in a public database; include a brief description of the data set or model with the number. If all data are in the paper and supplementary materials, include the sentence “All data are available in the main text or the supplementary materials.”

