## Supplementary Materials for "Huntingtin is an RNA-binding protein and participates in NEAT1-mediated paraspeckles"

Science

AAAS

**The PDF file includes:**

Materials and Methods
Supplementary Text
Figs. S1 to S9
References

### Materials and Methods

#### *Recombinant HTT-HAP40 Q23 Purification*

Expression of HTT-HAP40 Q23 protein was performed in Sf9 insect cells as previously described (*1–4*). Plasmids were transformed into DH10Bac E. coli competent cells, plated onto LB-agar plates, and then successfully transformed white colonies were selected and grown in 3 mL LB media supplemented with 50 µg/mL kanamycin, 7 µg/mL gentamicin, and 10 µg/mL tetracycline. Bacmids were purified using a Miniprep kit (Qiagen) but following cell lysate neutralization, the supernatants were mixed with 0.8 mL sterile isopropanol in a fresh tube and incubated on ice for 10 mins to precipitate the bacmid DNA. The DNA was pelleted by centrifugation, the supernatant removed, and the pellets left to airdry prior to resuspension in 50 µL of elution buffer from the kit. Exponentially growing Sf9 cells were diluted to 4 x 105 cells/mL in serum free insect media and 0.5 mL cell suspension/well used to seed 24-well plate to form a monolayer of cells. Plates were incubated at 27 °C for 1 h. Per well seeded in the 24 well plate, 2 µL of the X-tremeGENE 9 transfection reagent was mixed with 100 µL unsupplemented insect cell growth medium and 10 µL of a 0.2 µg/µL solution of recombinant bacmid DNA. The transfection mixture was incubated for 15-20 min to enable complex formation before addition of this mixture to a seeded well in the 24 well plate. Transfection plates were incubated for 4-5 h at 27 °C before addition of 1.5 mL of insect serum-free medium supplemented with 10% (v/v) of heat inactivated fetal bovine serum and 1% (v/v) antibiotic-antimycotic (100 units/mL of penicillin, 100 µg/mL of streptomycin, and 0.25 µg/mL of amphotericin B). Cells were incubated at 27 °C incubator for 72-96 h prior to harvesting of the P1 virus stocks and the recombinant virus titre was sequentially amplified by this method to generate P3 virus stocks.

Cells at ~4.5 million cells/ml were infected with 8 mL P3 recombinant baculovirus per 4 L of culture in 5 L reagent bottle and grown at 37°C until cell viability reached 80-85%, normally ~72 hours post-infection. The volume of culture grown can be scaled according to the required yield of protein. For full-length HTT-HAP40, a ratio of 1:1 HTT: HAP40 P3 baculovirus was used for infection.

For purification, cells were harvested by centrifugation (JLA8.1000 (Beckman), 2500 rpm, 10 °C, 10 min) and resuspended in ~40 mL lysis buffer (20 mM HEPES pH 7.4, 300 mM NaCl, 2.5% (v/v) glycerol) per L of production. Cell suspensions were lysed with two freeze-thaw cycles. Lysates were then diluted 3-fold in lysis buffer and clarified by centrifugation (JLA16.250 (Beckman), 14,000 rpm, 10 °C, 1 h). The supernatant was incubated with ~1 mL anti-FLAG resin (Sigma) per L of production processed and rocked for 2 h at 4 °C. The resin was washed with ~100 CV lysis buffer and then proteins were eluted with 5 CV lysis buffer supplemented with 250 µg/mL 3x FLAG peptide. Crude protein samples from FLAG-eluted fractions were then pooled, concentrated using Amicon spin filters to ~0.1 CV (MWCO 100,000) and purified by size-exclusion chromatography using a Superose 6 Increase 10/300 GL column (Cytiva Life Sciences) in buffer containing 20 mM HEPES pH 7.4, 300 mM NaCl, 2.5% (v/v) glycerol, 1 mM TCEP. To avoid overloading the gel filtration column, only 0.5-1 mL sample was applied per run. Peaks corresponding to monodisperse protein were pooled, concentrated and flash frozen at 5-10 mg/mL and stored at -80°C until use. Sample purity was assessed by SDS-PAGE.

***Surface Plasmon Resonance***

Studies were performed using a Biacore T200 (GE Health Sciences). An SA chip was primed with 3 x 60 s injection with 50 mM NaOH, followed by approximately 500 response units (RU) of biotinylated nucleic acid substrate diluted in assay buffer (20 mM HEPES pH 7.4, 50 mM KCl, 2 mM MgCl2, 1 mM TCEP, 2.5% glycerol (v/v), 0.005% Tween-20 (v/v)) coupling to flow channels, with an empty flow channel used for reference subtraction. Following substrate capture, assay buffer was flowed over the chip until a stable baseline was achieved. Protein dilutions were prepared in assay buffer and experiments were performed using multicycle kinetics with 600 s contact time, 600 s dissociation time and 30 µL/min flow rate at 20 °C. KD values were calculated using steady-state affinity fitting with the Biacore T200 evaluation software.

***Fluorescence Polarization Assay***

Experiments were performed in 384-well black polypropylene PCR plates (Axygen) in 20 µL volume. In each well, 18 µL protein solutions in assay buffer (20 mM HEPES pH 7.4, 50 mM KCl, 2 mM MgCl2, 1 mM TCEP, 2.5% glycerol (v/v), 0.005% Tween-20 (v/v)) were diluted, followed by the addition of 2 µL of 50-500 nM FITC-labelled nucleic acid substrate to each well. Following 1 min centrifugation at 1000 RPM, the plate was incubated for 10 minutes before FP measurements with a BioTek Synergy 4 (BioTek) at excitation and emission wavelengths of 485 and 528 nm, respectively. The data was processed in GraphPad Prism using Sigmoidal, 4PL, X is log(concentration) fit.

***Electrophoretic Mobility Shift Assay (EMSA)***

1% agarose gel was prepared with 2µL of ethidium bromide (BioShop Canada lnc., cat no. ETB444.1). HTT-HAP40 Q23 (WT) protein was serially diluted from 15µM to 0.2µM in the assay buffer (20 mM HEPES pH 7.4, 50 mM KCl, 2.5% glycerol (v/v), 2 mM MgCl2, 0.005% Tween-20 (v/v), 1 mM TCEP). The ssRNA and dsDNA oligonucleotide substrates were added in each reaction tube at a 1 µM final concentration and incubated on ice for 10 minutes. For NEAT1 fragments, the concentration used was 0.25 µM. The gel was run in 0.5X TAE buffer at 100V for 40 min for shorter oligos and 1 hr for NEAT1 fragments. The agarose gels were imaged under a UV transilluminator.

***Cell Culture***

Immortalised control and HD patient-derived fibroblasts were a kind gift from Professor Ray Truant and have been previously described (*5*). TruHD Q43/Q17, TruHD Q50/Q40, TruHD Q43/Q19, TruHD Q40/Q18 were derived from HD patients while TruHD Q19/Q17, TruHD Q21/Q17 were derived from a control subject. These cells were cultured in DMEM (Life Technologies #10370) with 15% (v/v) fetal bovine serum (FBS; Gibco #12484-028), 1X GlutaMAX (Life Technologies #35050) and 1x penicillin Streptomycin antibiotics. Cells were grown at 37°C with 5% CO2.

RPE1 cells (American Type Culture Collection) were cultured in 1:1 DMEM/Nutrient Mixture F-12 (DMEM/F12; Life Technologies #11330) with 10% (v/v) fetal bovine serum (FBS; Gibco #12484-028) and 0.01% hygromycin. Cells were grown at 37°C with 5% CO2.

Human embryonic kidney line 293 (HEK293T) wild type and HTT null cells were unauthenticated and a kind gift from the laboratory of Marcy MacDonald (*6*). These cells were cultured in DMEM (Life Technologies #10370) with 10% (v/v) fetal bovine serum (FBS; Gibco #12484-028), 1X GlutaMAX (Life Technologies #35050) and 1x penicillin Streptomycin antibiotics. Cells were grown at 37°C with 5% CO2.

***Isogenic NPC Differentiation***

hESCs were plated at a density of 30,000-50,000/cm2 per well in NPC+ media [equal parts DMEM/F12 (Thermo Fisher Cat No. A4192001) and NeuroBasal medium (Thermo Fisher Cat No. A3582901); 50µg/ml BSA (Thermo Fisher Cat No. B14); 1X PenStrep (Thermo Fisher Cat No. 15140122); 1X MEM NEAA (Thermo Fisher Cat No. 11140050); 1X N2 (Thermo Fisher Cat No. 17502001); 1X B27 without Vit A (Thermo Fisher Cat No. 12587010); 10µg/ml human LIF (Merck Millipore Cat No. LIF1010); 2µM SB431542 (Stem Cell Tech Cat No. 72232); 3µM CHIR99021 (Stem Cell Tech Cat No. 72052); 0.1µM Compound E (Stem Cell Tech Cat No. 73952)] supplemented with 10µM Y-27632 (Stem Cell Tech Cat No. 72304) on Geltrex (Thermo Fisher cat No. A1413301) coated plates. This is day 1 of differentiation. NPC+ medium was replaced on a daily basis till day 7. On day 7, the cells were passaged using Accutase (Thermo Fisher Cat No. A1110501), split at a ratio of 1:10 and plated in NPC- medium (NPC+ medium without Compound E) with 10µM Y-27632. The cells were passaged at confluency >90%. NPCs at passage 5 or higher were then subsequently used for experiments.

***UV-crosslinking***

10x106 cells were plated on 100mm culture plates. After 24hrs, the plates were washed with 1X PBS. Thereafter, 2ml of PBS to each of the plates and placed on a tray of ice. The whole setup was plated inside the UV crosslinker (VWR Cat No. 89131-484) without the lid. The plate was placed 15cm from the light source. The crosslinker was operated at 254-nm UV with an energy setting of 400mJoules/cm2 for 10 minutes. Post crosslinking, the cells were collected with a cell scraper and centrifuged at 4oC, 300g for 5 minutes. The supernatant was discarded, and the pellets flash frozen in liquid nitrogen and stored at -80oC freezer.

***RNA-immunoprecipitation from isogenic NPCs and Fibroblasts***

The pellets were retrieved from -80°C and immediately resuspended in 1mL of cold CLIP lysis buffer (50mM Tris-HCl pH 7.4, 100mM NaCl, 1% NP-40, 0.1% SDS, 0.5% sodium deoxycholate, 1X protease inhibitor, and RNasin plus ribonuclease inhibitor). The cells were lysed on ice for 15 min and sonicated in Bioruptor at “low” setting, 4°C for 5 min with 30 sec on and 30 sec off. 4 Units of DNase were added, and the tubes were incubated in a thermomixer at 37°C with shaking at 1200 rpm for 30 min. Next, the tubes were transferred on ice and immediately 50mM final concentration of EDTA was added. The tubes were centrifuged at 20,000xg at 4°C for 15 min to pellet debris. The supernatant was transferred to a new tube.

125uL beads for each sample was washed 2X in 500µL cold CLIP lysis buffer, the beads were resuspended in 100uL cold CLIP lysis buffer and 10µg of EPR5526 (Abcam) anti-HTT antibody was added to the resuspended magnetic beads. The beads were rotated at 4°C for 1-2 hours. The tubes were centrifuged gently and placed on a magnetic rack; the supernatant was removed. The beads were further washed 2x in 500µL cold CLIP lysis buffer. Beads for control IP (without any antibody) were prepared in the same way. 50 µL of the cell lysates were kept for western blot input. The remaining cell lysates were added to the prepared magnetic beads and the tubes were kept rotating at 4°C overnight. Next day, the tubes were placed in a magnetic rack and the cell lysate flowthrough was collected. Beads were washed further 2x with 900 µL cold high salt wash buffer (50mM Tris-HCl pH 7.4, 500mM NaCl, 1mM EDTA, 1% NP-40, 0.1% SDS, 0.5% sodium deoxycholate) and 1x with a 500µL cold CLIP lysis buffer. 150uL of Proteinase K buffer (50mM Tris-HCl pH 7.4, 150mM NaCl, 1mM MgCl2, 0.05% NP-40, 1% sodium deoxycholate, and 1.2mg/mL Proteinase K) was added to the beads and incubated at 55°C for 30 min while shaking to digest proteins. After this reverse cross-linking step, the tubes were gently centrifuged using a table-top centrifuge and placed on a magnetic rack. The supernatants were transferred to new tubes and 600µL TRIzol reagent was added, and the tubes were incubated for 10 min at room temperature. The RNA was further purified using Direct-Zol RNA miniprep plus kit (Zymo Research, Cat. no. R2071). The purified RNA was eluted in RNases and DNase free water, and concentrations were estimated by nanodrop prior to library preparation.

***SDS-PAGE and western blotting for IP validation***

50 µg of protein lysate was used per sample. Samples were prepared using 4x NuPAGE LDS Sample Buffer (ThermoFisher Scientific; catalogue #NP0007) and 10x NuPAGE Sample Reducing Agent (ThermoFisher Scientific; catalogue #NP0004) and denaturing the samples at 70°C for 10 minutes. The denatured samples were electrophoresed at 120 volts for 3 hours on NuPAGE 4-12% Bis-Tris Proteins Gels (ThermoFisher Scientific; catalogue # NP0321BOX) in NuPAGE MOPS SDS Running Buffer (ThermoFisher Scientific; catalogue #NP0001). Samples were run in parallel with Full range rainbow MW marker (ThermoFisher Scientific, catalogue #RPN800E) and/or HiMark Pre-stained Protein Standard (ThermoFisher Scientific, catalogue #LC5699). Gels were wet-tank transferred to PVDF Western Blotting Membranes (Sigma-Aldrich, Cat #3010040001; activated in 100% methanol for 1-2 minutes prior to use) in tris-glycine (with 10-20% methanol) overnight (16-24 hours typically) at 4°C using a constant voltage of 30V. The next day, membranes were blocked in 5-10% w/v milk dissolved in 1xTBS + 0.1% Tween-20 (TBST) for 1 hour at room temperature. Blots are then incubated with primary antibody at room temperature for 2 hours using the same solution used for blocking, washed 3 times in 1xTBST at room temperature (10 minutes/wash), incubated with secondary antibody at room temperature for 1 hour in the same solution used for blocking, washed 3 times in TBST at room temperature (10 minutes/wash), and then detected with ECL (GE Healthcare Amersham ECL™ Prime Western Blotting Detection Reagent, Cat #RPN2232) by autoradiograph. Densitometric quantification of bands is performed using Image Studio Lite Version 5.2 (LI-COR Biosciences).

For HTT immunoprecipitation western blot from fibroblasts:

Primary antibody: Anti-HTT Clone 1HU 4C8 (1:1000, EMDmillipore, Catalogue #MAB2166)
Secondary antibody: Peroxidase-AffiniPure Sheep Anti-Mouse IgG H+L (1:2000, Cedarlane Labs, catalogue #515035062)
For HTT immunoprecipitation western blot from NPCs:
Primary antibody: Anti-HTT 2B7 (2:1000, CHDI-90000830-5, #CH03023)
Secondary antibody: Anti-mouse IRDye 800CW (1:10000, LI-COR Biosciences, 926-32210 or 926-32212).

***RNA Library Preparation***

RIP-Seq libraries were prepared by using SMARTer Stranded Total RNA-Seq Kit version 2 (Pico Input Mammalian; Takara, 634413) as per the manufacturer's instructions. Subsequently, the libraries were subjected to paired-end sequencing with a read length of 150 bp on the Illumina HiSeq X Ten platform, ensuring a minimum sequencing depth of 30 million reads per sample. For both input and immunoprecipitated (IP) samples, cDNA synthesis was carried out utilizing the High-Capacity cDNA Reverse Transcription Kit (ThermoFisher, 4368814). Quantitative PCR (qPCR) was performed using the PowerUp SYBR Green Master Mix (Applied Biosystems, Cat. #A25742) on either the StepOnePlus Real-Time PCR System (Applied Biosystems) or the CFX96 Touch Real-Time PCR Detection System (Bio-Rad). To normalize the qPCR data, either U6, RPS28, and GAPDH was employed as the endogenous control gene. The fold change was calculated utilizing the ΔΔCt method.

***High Throughput Sequencing Data Alignment and Analysis***

RIP-seq reads were aligned to the human reference genome hg38 by using STAR (version 2.6.1c) (*7*) with the reference annotation GENCODE version 25 (*8*). The genes bound by HTT were defined as genes enrichment fold change (log2(IP/INPUT)) greater than 1 and P-adj < 0.05. Functional enrichment analysis was performed through the web server g:Profiler (*9*) using annotated genes as background.

***MEME motif analysis and sequence annotation***

For meme motif sequence analysis, the protocol used was followed as in Owens et al. (*10*) with modifications. Briefly, with the resulting human genome (hg38) mapped reads as bam files, peaks were called using MACS2 with a p-value threshold of 0.01. The input and IP reads were submitted jointly to account for background noise. The peaks were written out in BED format specifying the location of each peak in the human genome. 2000 peaks were sampled using BEDTools (2.30.0) and flanking regions were extracted from the sampled peaks. Sequences of sampled peaks and flanking regions were retrieved, and a background file was generated using fasta-get-markov -m 0. A de novo motif file was generated using MEME (4.5.1) with options -rna -nmotifs 20 -w 20 -maxsize 1000000 -mod zoops.

***Reverse Transcription qPCR (RT-qPCR)***

For RT-qPCR, cDNA synthesis was carried out utilizing the High-Capacity cDNA Reverse Transcription Kit (ThermoFisher, 4368814). Quantitative PCR (qPCR) was performed using the PowerUp SYBR Green Master Mix (Applied Biosystems, Cat. #A25742) on either the StepOnePlus Real-Time PCR System (Applied Biosystems) or the CFX96 Touch Real-Time PCR

Detection System (Bio-Rad). To normalize the qPCR data, either U6, RPS28, and GAPDH was employed as the endogenous control gene. The fold change was calculated utilizing the ΔΔCt method.

***RNA Preparation from Patient Brain Tissues***

Tissues (stored and -80°C and kept immersed liquid nitrogen during handling) were crushed with a frozen metal mortar and pestle partway buried in dry ice, and frozen crushed tissues were immediately transferred to a 1.4 mm Acid Washed tube pre-filled with Zirconium Beads and 300-1000 µL of TRIzol reagent. Smaller tissues were directly inserted into tubes without crushing. Tubes were inverted to ensure immersion of the whole tissue and were placed at room temperature for 10 minutes to allow the TRIzol reagent to denature and remove proteins bound to RNA. Tubes were placed on ice after ten minutes and then placed in a MagNA Lyser Instrument (Roche; item #03358968001). Tubes were oscillated at 7000 OSC 3 times for 20 seconds each oscillation, with a 3-minute incubation on ice between each 20 second oscillation. TRIzol was transferred to a different tube, RNA precipitated by an equal volume of 100% EtOH and then purified using the Direct-zol RNA purification kit using the manufacturers protocol, which includes in-column DNase treatment (Zymo research; catalog # R2071). Whole RNA was reverse transcribed using the SuperScript IV First-Strand Synthesis System kit using the manufacturers protocol (ThermoFisher Scientific; catalogue #18091050).

***HTT Knockdown Using SiRNAs***

siRNAs for HTT knockdown were purchased from Horizon Discovery (#M-003737-02-0005) and control siRNA were purchased from Sigma (#SIC001-5X1NMOL). 2.5 × 105 cells were seeded per 2mL of media per well in a 6-well tissue culture plate. The cells were transfected at seeding by Lipofectamine RNAiMAX Transfection Reagent (Thermo Fisher #13778150), per manufacturer protocol. For every well in 6-well plates: 150 µL of OptiMEM Medium (Gibco #31985070) was mixed with 9 µL of RNAiMAX reagent and 150 µL of Opti-MEM was mixed with 3 µL of 10 µM siRNA, separately. The two mixtures were then mixed to form a 300 µL mixture and incubated at room temperature for 5 min to allow the formation of siRNA−lipid complexes. The siRNA−lipid complex mixture was added to the wells at 250 µL/well for each 6-well plate.

***Cell Lines for HTT Knockdown by ShRNAs (RPE1 And Fibroblasts)***

HTT shRNA pNT153 and pNT154 from TRCN, and negative control shGFP from Dr Hansen He's lab, sequences listed below, were cloned into lentiviral vector Tet-pLKO-neo (Addgene #21916) using Dr. Dmitri Wiederschain's protocol (ref. The "all-in-one" system for the inducible expression of shRNA, Novartis Developmental and Molecular Pathways, Cambridge, MA, USA). To produce lentivirus, shRNA and packaging plasmids were cotransfected into HEK293 cells using X-tremeGene HP DNA Transfection Reagent (Millipore, catalogue 6366236001). Lentivirus was collected 48 hours later and filtered through 0.45 um acrodisc filter.

Tru-HD-Q21/Q18, Tru-HD-Q43/Q17 and RPE1 cell lines were infected with virus along with polybrene (8ug/ml) for 24 hours then the cells were growing in regular medium for 24-48 hours

before screening with neomycin. For Tru-HD-Q21 and Q43, 400 ug/ml of neomycin was used; for RPE1, 800 ug/ml neomycin was used. It took 3 weeks to obtain the stable cell lines.
pNT153: TRCN0000323037
Hairpin Sequence: 5'-CCGG-GCACTCAAGAAGGACACAATA-CTCGAG-TATTGTGTCCTTCTTGAGTGC-TTTTTG-3'
pNT154: TRCN0000350710
Hairpin Sequence: 5'-CCGG-TGGTTCAGTTACGGGTTAATT-CTCGAG-AATTAACCCGTAACTGAACCA-TTTTTG-3'
shGFP-Hairpin sequence: 5'- CCGG-CCACATGAAGCAGCACGACTT-CTCGAGAAGTCGTGCTGCTTCATGTGG-TTTTTG-3'

***SDS-PAGE and Western blot analysis***

Fibroblast and RPE1 cells after siRNA or shRNA HTT knockdown were scraped and centrifuged at 4°C at 1500 rpm. Cell pellets were lysed in a lysis buffer (20mM Tris pH 8.0, 150mM NaCl, 10mM MgCl2, 1mM EDTA, 0.5% Triton X-100, 1X protease inhibitor, benzonase and 1% SDS) for 10 min on ice and centrifuged at 14,000 rpm for 5 min. Supernatant was collected and total protein concentration was estimated using the Pierce BCA protein assay kit (Thermo Scientific #23225). Equal concentrations of total protein were loaded into a precast NuPAGE 4-12% Bis-Tris gel (Invitrogen #NP0323BOX). Proteins were separated by SDS–PAGE in 1X MOPS buffer (Life Technologies #NP0001) for 3hr at 120 volts, and electroblotted onto 0.2μm polyvinyl difluoride (PVDF) blotting membrane (Amersham Hybond #GE10600021) overnight at 30V in 1x Transfer buffer (25 mM Tris, 192 mM Glycine, 0.375% SDS, and 10% methanol). Next day, the membrane was blocked in 5% (w/v) skimmed milk prepared in 1xPBST buffer and incubated with primary antibodies EPR5526 (2:10,000; Abcam ab109115) or 2B7 (CHDI-90000830-5, #CH03023) and/or anti-β-actin (3:10,000; Santa Cruz sc-47778) overnight at 4°C. The membranes were further washed 3x for 10 min with 1xPBST and incubated with secondary antibodies (anti-rabbit, 926-32211, IRDye 800CW, LI-COR Biosciences, 1:10 000; anti-mouse IRDye 800CW, LI-COR Biosciences, 926-32210 or 926-32212, 1:10000) in blocking buffer for 1h at room temperature. Membranes were washed 3x for 10 min with 1xPBST and imaged on a LI-COR Odyssey® CLx.

***RNA FISH and immunofluorescence (IF)***

Cells were grown on 18mm round #1.5 coverglass in a 12-well glass-bottom cell culture dishes to approximately 80% confluence prior to fixing with 4% paraformaldehyde (PFA) solution in 1xPBS. Next day, the growth medium was removed, and the cells were washed with 1XPBS. The cells were then fixed with 4% PFA and incubated for 10 minutes at room temperature. The 4% PFA was removed, and the fixed cells were washed 3x with 1xPBS. Stellaris FISH probes to the NEAT1 5'-region (LGC Biosearch Technologies, SMF-2036-1) and NEAT1 middle-region (LGC Biosearch Technologies, SMF-2037-1) and the FISH was performed according to the manufacturer's instructions and as in Yamazaki et al. (*11*). The fixed cells were immersed in 70% ethanol for at least 1 h at 4°C for permeabilization. After removing the 70% ethanol, wash buffer (2X SSC and 10% formamide) was added and the coverslips were incubated for 5 min. Then, the

50 μL hybridization solutions (2X SSC, 100 mg/ml dextran sulfate, and 10% formamide) containing Stellaris NEAT1 probes (final concentration: 125 nM) were dropped onto the coverslips in the humidified chamber and were incubated for 16 h in the dark. The coverslips were washed with wash buffer at 37°C in the dark for 30 min and with 2X SSC at room temperature for 5 min. For just FISH, the coverslips were mounted with VECTASHIELD Hard Set Mounting Medium with DAPI (Vector).

The coverslips were subsequently washed with 1XPBS and incubated with blocking solution (Intercept blocking buffer, LI-COR, # 927-70001 and 1XPBST) for blocking at room temperature for 1 h. Then, the coverslips were incubated with HTT primary antibodies (N17-phospho, 1:250) in blocking solution at room temperature for 1 h, washed three times with 1XPBS for 5 min, incubated with secondary antibody anti-rabbit (Alexa Fluor 488) in blocking solution at room temperature for 30 min, and washed 3x with 1XPBS for 5 min. The coverslips were mounted with VECTASHIELD Hard Set Mounting Medium with DAPI (Vector). Confocal images were acquired using a confocal microscope (Leica SP8).

***RNA stability assay***

For RNA stability assay, the protocol was followed as in Ratnadiwakara et al. (*12*) with modifications. 1x105 RPE1 cells containing shGFP and HTT shRNA plasmids were seeded per well in a 6-well plate for 5 timepoints (t=0 to t=4 hours). The plate was incubated overnight at 37ºC, and 2ug/mL doxycycline was added in each well on the next day for initiating HTT knockdown. On the fifth day, cells from t=0 timepoint were collected, spin at 470xg for 3min at 20C, and frozen in -80C. To the remaining wells, 30 uL of 1mg/mL Actinomycin D was added to obtain a final concentration of 10ug/mL in 3 mL of culture media. The samples were collected at 1-, 2- and 4-hour time points and the pellet was frozen in -80C until use. The RNA was isolated using the Qiagen RNA miniprep kit and RT-qPCR was performed. The fold change was calculated utilizing the ΔΔCt method and all samples were normalized to the control t=0 timepoint.

***Image processing and NEAT1-paraspeckle count***

The z-stack images were acquired using Leica TCS SP8 confocal microscope (Leica DMi8 CS inverted stand) with 405 nm, 552 nm and 488 nm lasers. The images were processed and analyzed in LASX office software (version 1.4.1) and ImageJ. The NEAT1-paraspeckles number and intensities were quantified in over 150-200 cells over three trials using an open-source speckle counting pipeline in CellProfiler (version 4.2.6).

***Data visualization and analysis***

Data analysis and visualization were conducted using R (version 4.0.0), packages used: ggplot2_3.4.1, clusterProfiler_3.16.1, ComplexUpset_1.3.3 and venn_1.11, and GraphPad Prism (version 10.1.1). The outcomes of all statistical tests including P values and number of samples are included in the figure panels or the corresponding figure legends. Significance was defined as any statistical outcome that resulted in a P value of less than 0.05, unless otherwise indicated.

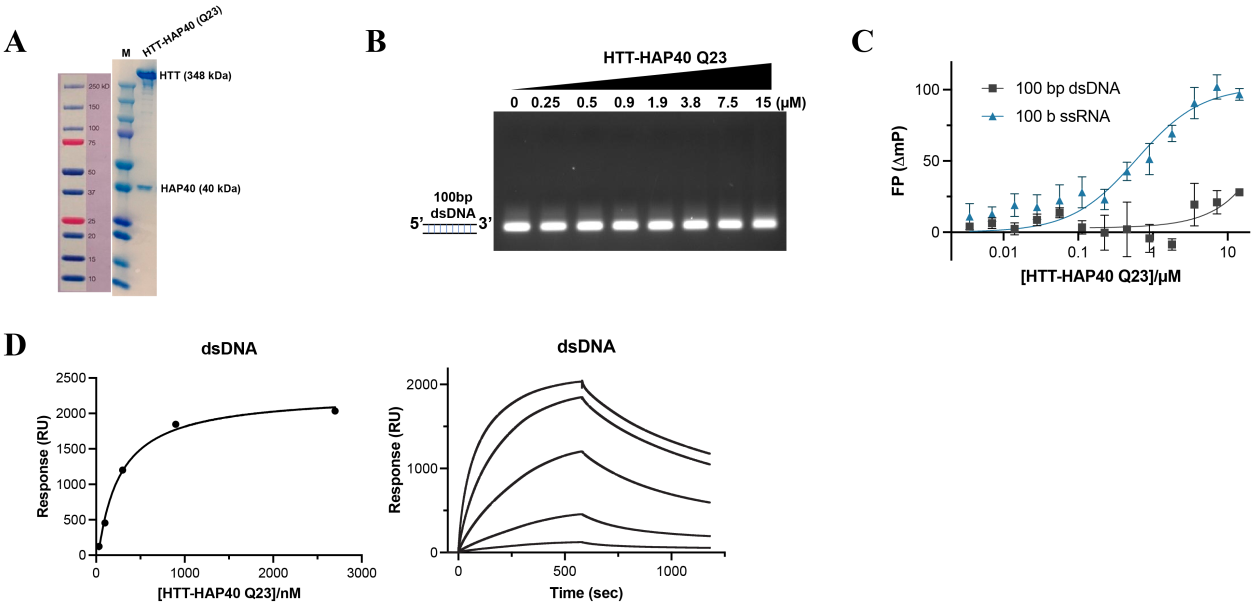


**Fig. S1**. **HTT-HAP40 prefers to bind RNA and prefers G-rich sequences. (A)** SDS-PAGE analysis showing recombinantly purified HTT-HAP40 Q23 protein after gel filtration chromatography purification. **(B)** Representative EMSA image of increasing HTT-HAP40 Q23 protein (0–15 μM) binding with 1 μM of 100bp dsDNA. DNA is in black. EMSA, electrophoretic mobility shift assay. **(C)** Representative FP binding curve of HTT-HAP40 Q23 and 100-mer random ssRNA ($K_D$ = 1 ± 0.6 μM), dsDNA ($K_D$ = NC). NC, not calculated, outside of range of protein concentrations tested. **(D)** Representative SPR binding curve and sensorgram of HTT-HAP40 Q23 and 100bp dsDNA ($K_D$ = 2.0 ± 0.2 μM).

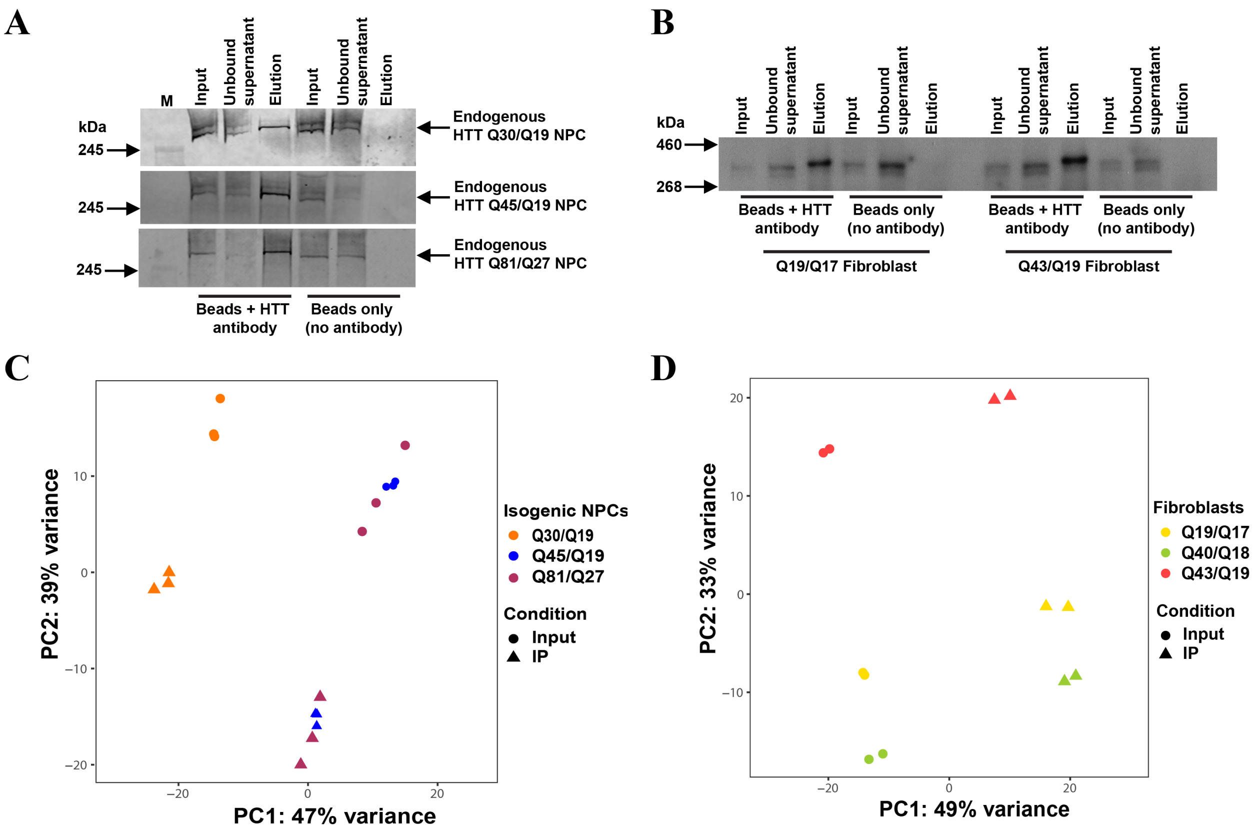


**Fig. S2. HTT RIP-seq analysis in isogenic NPCs and fibroblasts. (A-B)** Validation of HTT IP protocol through western blot analysis in NPCs (A) and fibroblast cell lines (B). **(C-D)** Principal components analysis (PCA) of RIP-seq dataset of WT and HD isogenic NPCs (C) in triplicates, and fibroblast (D) cell lines in duplicates.

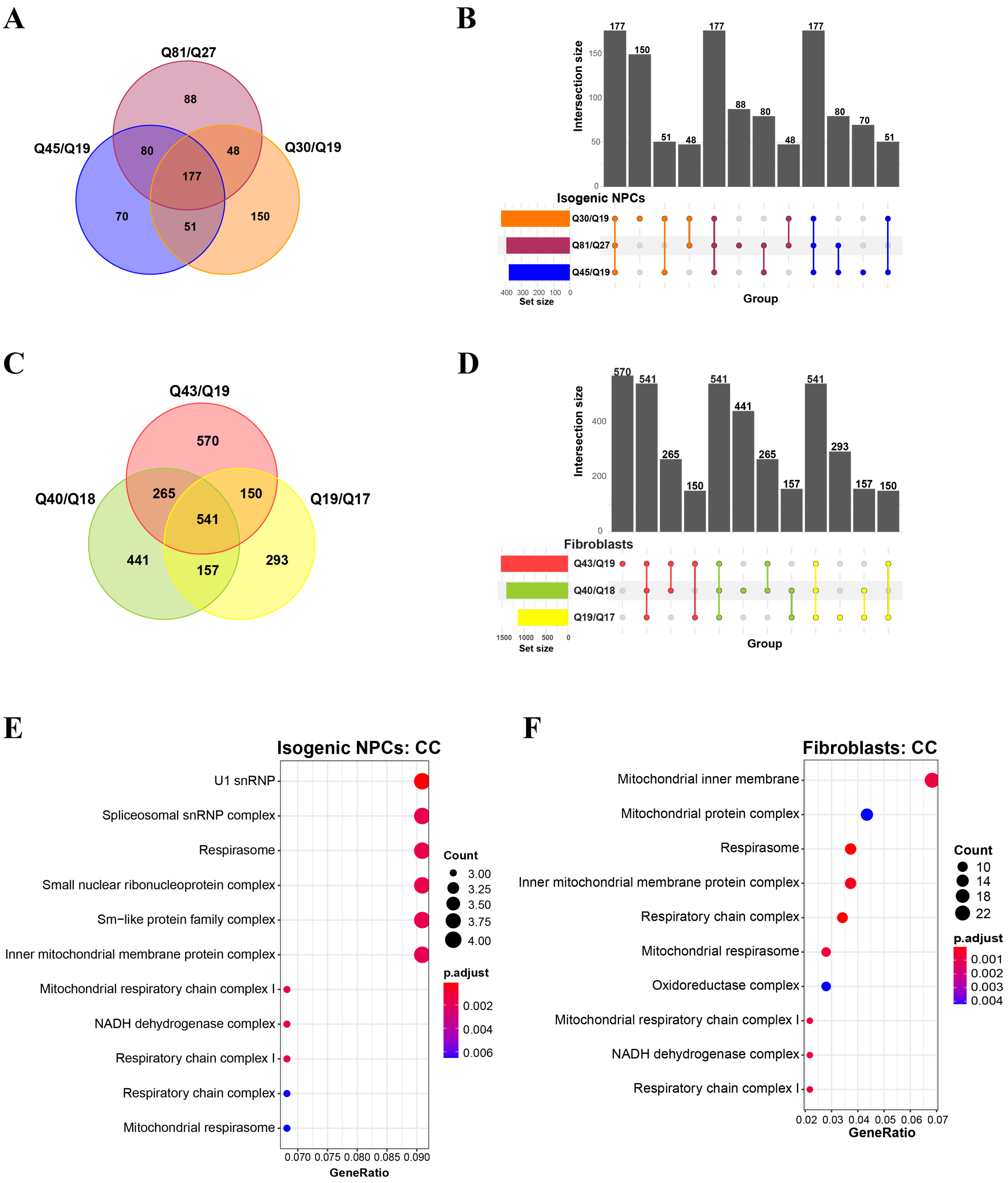


**Fig. S3. HTT RIP-seq analysis in isogenic NPCs and fibroblasts. (A)** Venn diagram representing enriched RNA transcripts from IP samples of WT and HD NPCs. **(B)** Bar graph representing unique and overlapped enriched RNA transcripts from NPCs IP samples. **(C)** Venn diagram representing enriched RNA transcripts from IP samples of WT and HD fibroblasts IP samples. **(D)** Bar graph representing unique and overlapped enriched RNA transcripts from fibroblasts IP samples. **(E-F)** GO enrichment analysis for cellular component (CC) in WT and expanded NPCs (E) and fibroblasts (F).

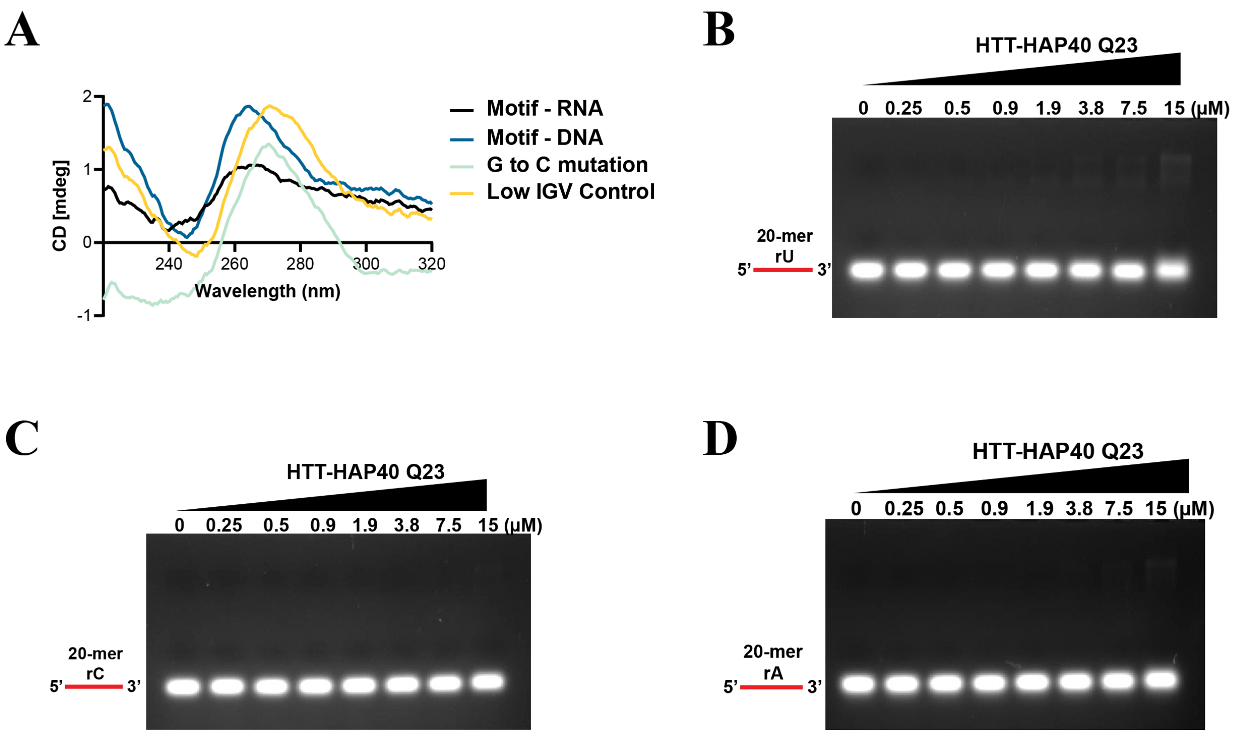


**Fig. S4. (A)** Circular Dichroism (CD) spectroscopy analysis for secondary structure folding of identified RNA motif (Motif-RNA), RNA to DNA motif (Motif-DNA), G to C mutated RNA, and low IGV RNA. **(B-D)** Representative EMSA images of increasing HTT-HAP40 Q23 protein (0–15 µM) binding with 1 µM of indicated 20-mer rU (B), rC (C), and rA (D) substrates. RNA is in red. EMSA, electrophoretic mobility shift assay.

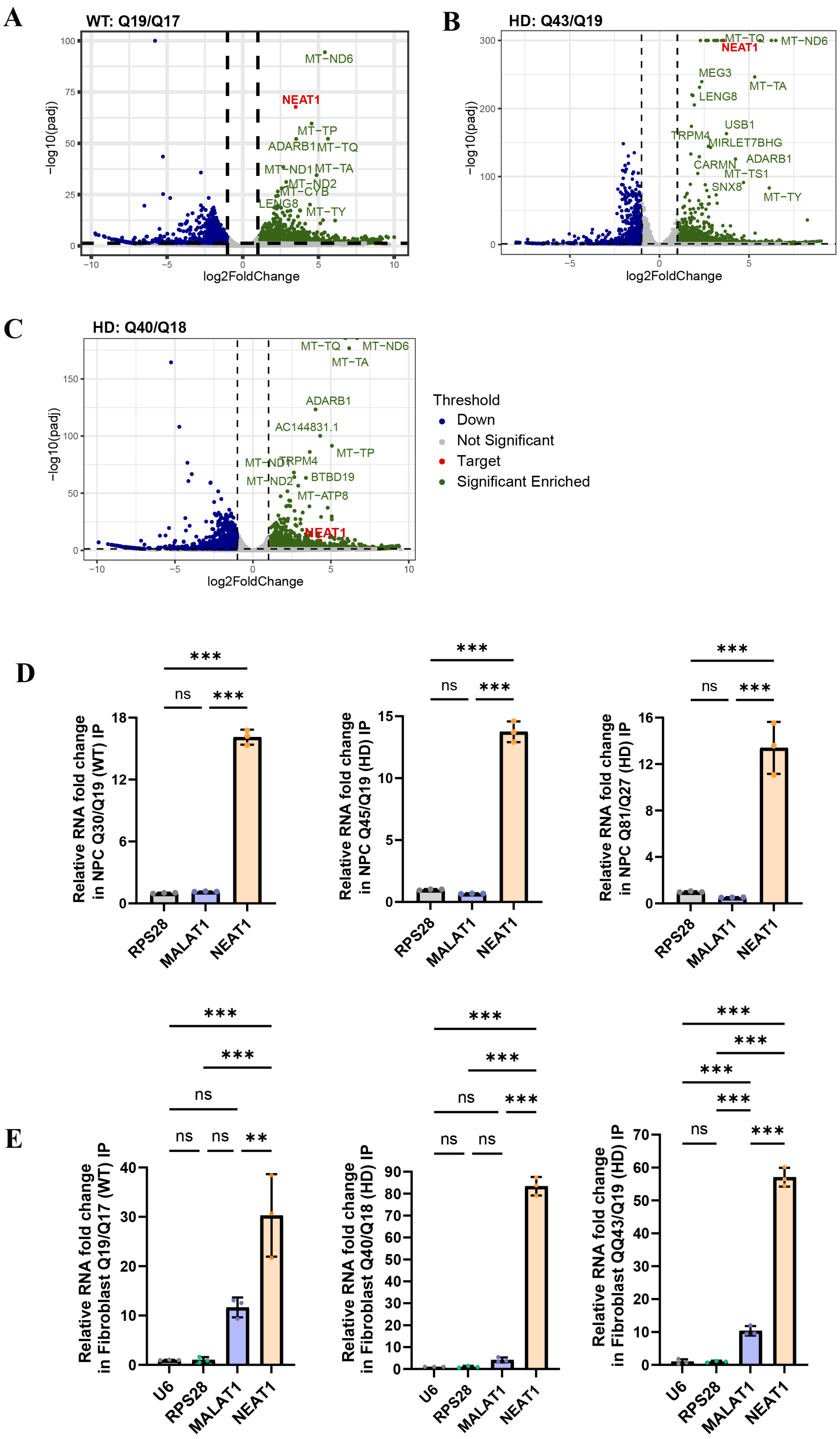
A
WT: Q19/Q17
B
HD: Q43/Q19
C
HD: Q40/Q18
−log10(padj)
log2FoldChange
MT−ND6
NEAT1
MT−TP
ADARB1
MT−TQ
MT−ND1
MT−TA
MT−ND2
MT−CYB
LENG8
MT−TY
MEG3
USB1
TRPM4
MIRLET7BHG
CARMN
MT−TS1
SNX8
AC144831.1
BTBD19
MT−ATP8
Threshold
Down
Not Significant
Target
Significant Enriched
D
Relative RNA fold change in NPC Q30/Q19 (WT) IP
Relative RNA fold change in NPC Q45/Q19 (HD) IP
Relative RNA fold change in NPC Q81/Q27 (HD) IP
RPS28
MALAT1
NEAT1
ns
***
**
E
Relative RNA fold change in Fibroblast Q19/Q17 (WT) IP
Relative RNA fold change in Fibroblast Q40/Q18 (HD) IP
Relative RNA fold change in Fibroblast QQ43/Q19 (HD) IP
U6

**Fig. S5. LncRNA NEAT1 enrichment validation using RT-qPCR. (A-C)** NEAT1 is significantly enriched in the WT and HD IP samples derived from fibroblast cells ($Log_2$ fold change cutoff: 1; p-value cutoff: 0.05). **(D-E)** Both form-specific (total NEAT1) and long form-specific (NEAT1_2) enrichment evaluation by RT-qPCR from WT and expanded isogenic NPCs (D) and fibroblasts (E). Three abundant coding and noncoding transcripts (RPS28, U6 and MALAT1) are shown as negative controls. Data was analyzed using one-way ANOVA. Data are shown as mean ± s.d.; $n = 3$. ***P < 0.001, **P < 0.01, ns, not significant.

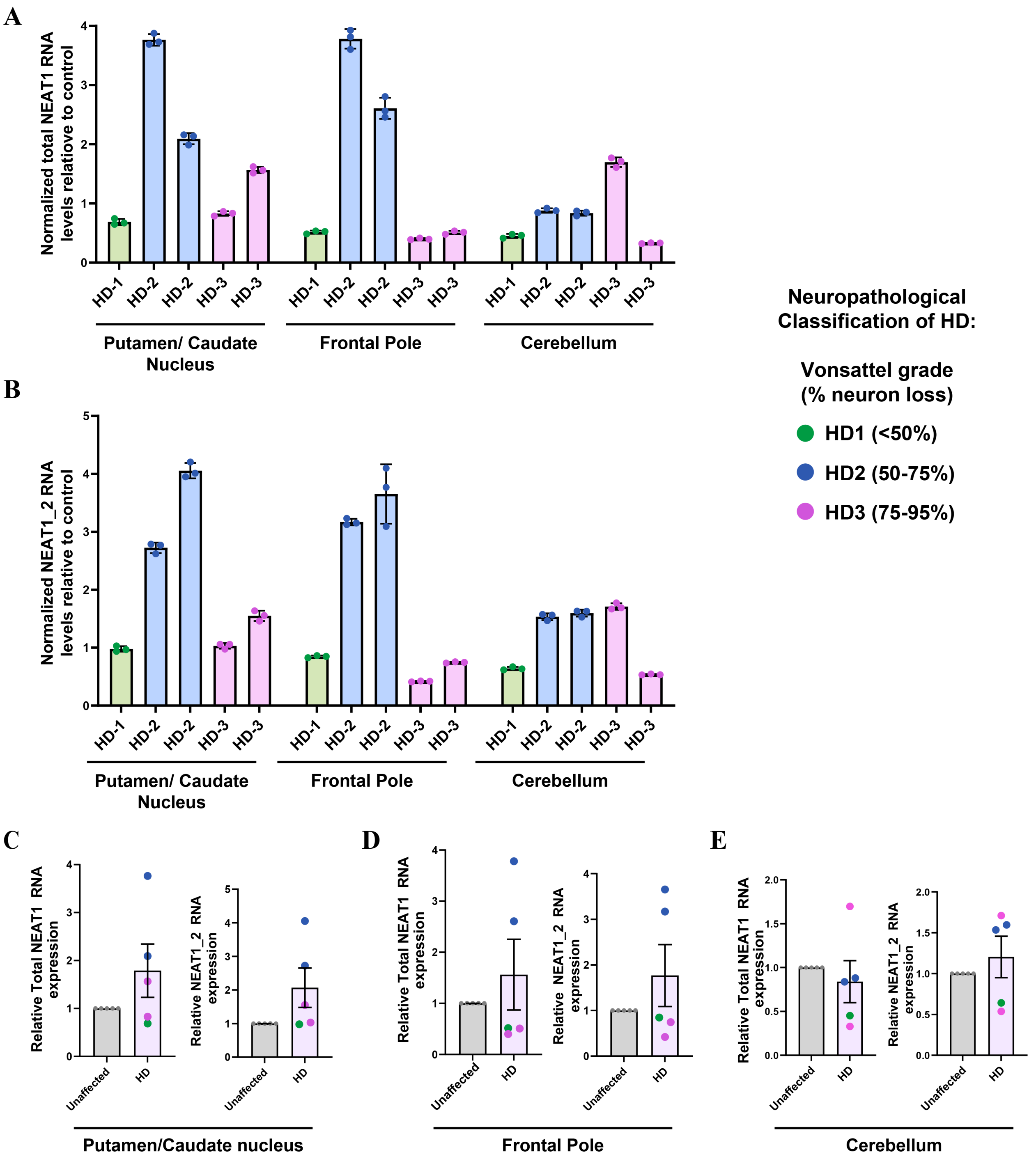


**Fig. S6. Altered lncRNA NEAT1 levels in HD patient brain tissues (A-B)** Total NEAT1 (A) and long NEAT1 (NEAT1_2) (B) expression levels in putamen, frontal pole, and cerebellum regions of human HD patient brain at different HD-grades. The HD patient tissues were normalized to age and sex matched unaffected brain tissues for each region, *n* = 5 HD and unaffected individuals/group/tissue, 3 technical replicates/person. **(C-E)** Both isoform specific NEAT1 (total NEAT1) and NEAT1_2 expression levels in putamen, frontal pole, and cerebellum regions of human HD patient brain compared to age and sex matched unaffected brain tissues (gray) for each region. *n* = 5 HD and unaffected individuals/group/tissue, average of 3 technical replicates/person. Dots indicate striatal neuropathological grade (HD 1–3). U6 was used as a control gene. No statistics applied due to limited patient sample size.

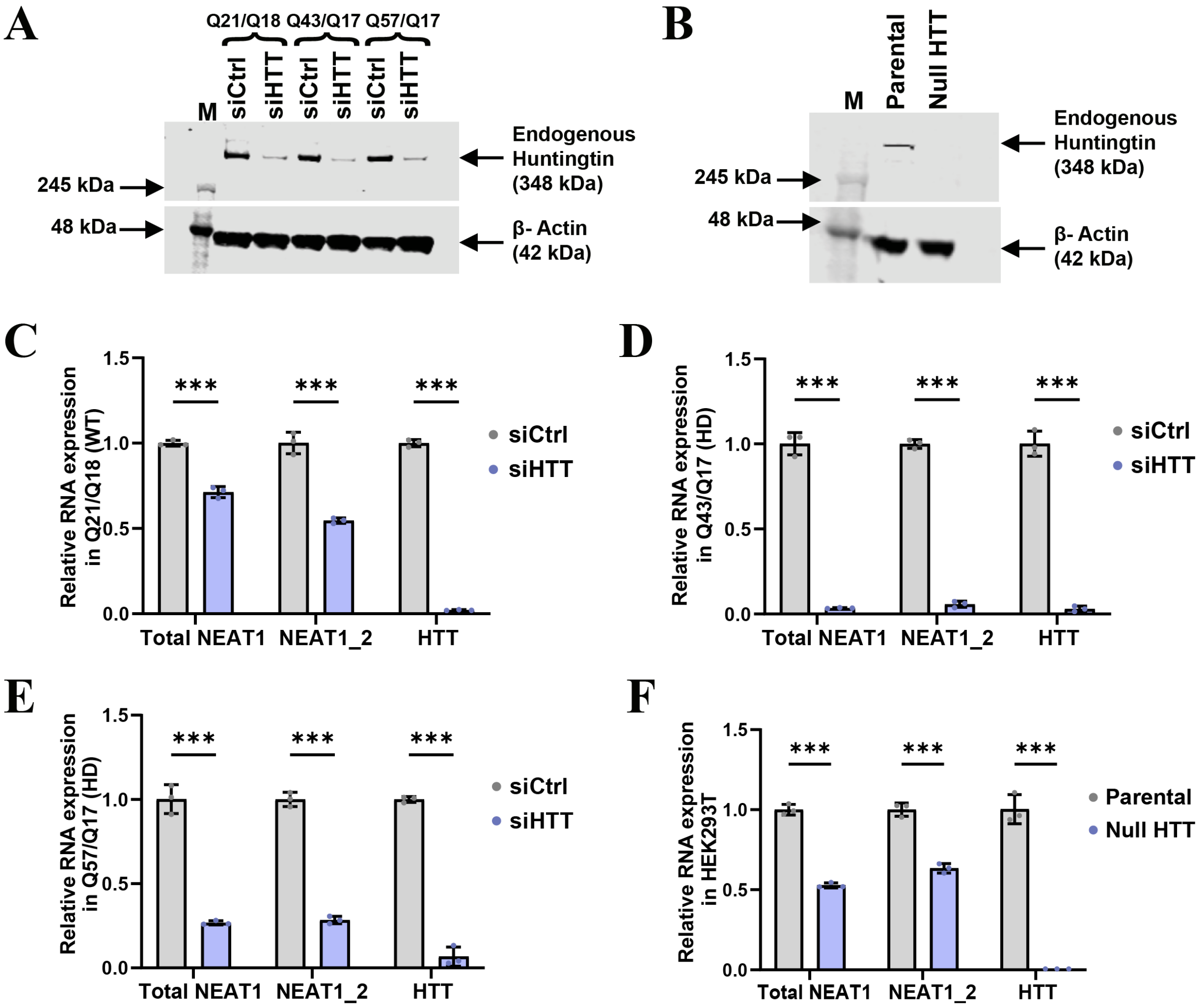


**Fig. S7. HTT knockdown reduces lncRNA NEAT1 levels. (A)** Western blot analysis of HTT protein knockdown by siRNAs in WT (Q21Q18), HD (Q43Q17) and HD (Q57Q17) fibroblast cell lines. **(B)** Western blot analysis of HTT expression in parental and HTT knockout HEK293T cells. **(C-E)** RT-qPCR quantification of levels of lncRNA NEAT1 isoforms upon HTT knockdown by siRNAs in WT (Q21Q18), HD (Q43Q17) and HD (Q57Q17) fibroblasts, respectively. **(F)** RT-qPCR quantification of levels of lncRNA NEAT1 isoforms in HEK293T parental and Null HTT cell lines. U6 was used as control gene and data were analyzed using 2-way ANOVA. Data are shown as mean ± s.d.; n = 3. ***P < 0.001.

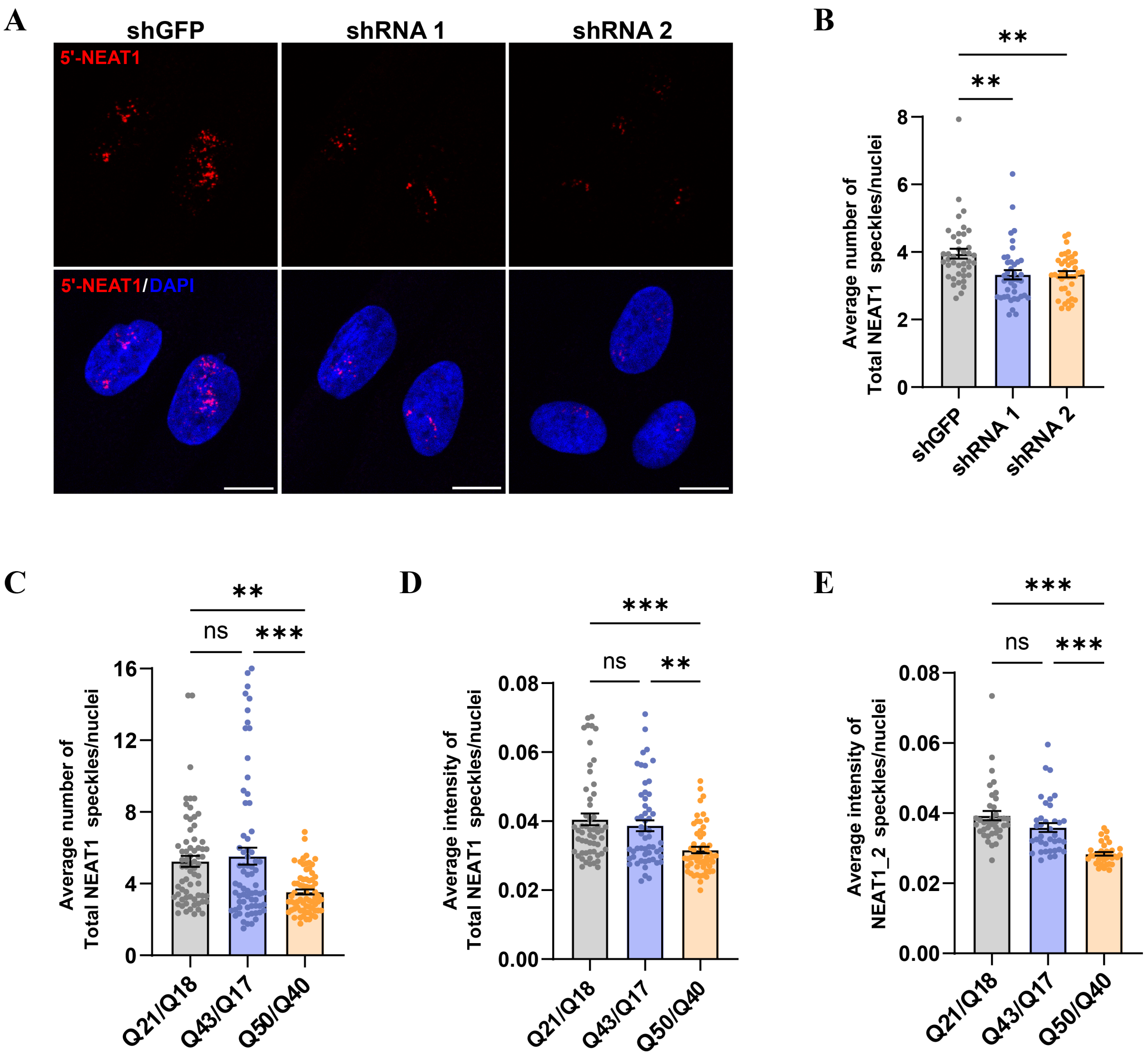


**Fig. S8. NEAT1 lncRNA levels reduce after HTT knockdown (A-B)** Representative images **(A)** and quantification **(B)** of NEAT1_1S positive foci in Q21Q18 fibroblast cell lines after HTT knockdown, assessed by FISH. Scale bar = 10 µm. For statistical analysis of the NEAT1_1S RNA foci, CellProfiler software was used, and images were processed using ImageJ (Fiji app). Data represent mean ± sem. Data were analyzed by ordinary one-way ANOVA with Tukey's test for multiple comparisons. ∗∗∗P < 0.001. **(C)** Quantification of average number of NEAT1_S foci in WT and HD fibroblast cell lines. **(D-E)** Quantification of average intensity of NEAT1_S and NEAT1_2L foci in WT and HD fibroblast cell lines. Data represents mean ± sem. Data were analyzed by ordinary one-way ANOVA with multiple comparisons. ∗∗∗P < 0.001, ∗∗P < 0.01 and ns, not significant.

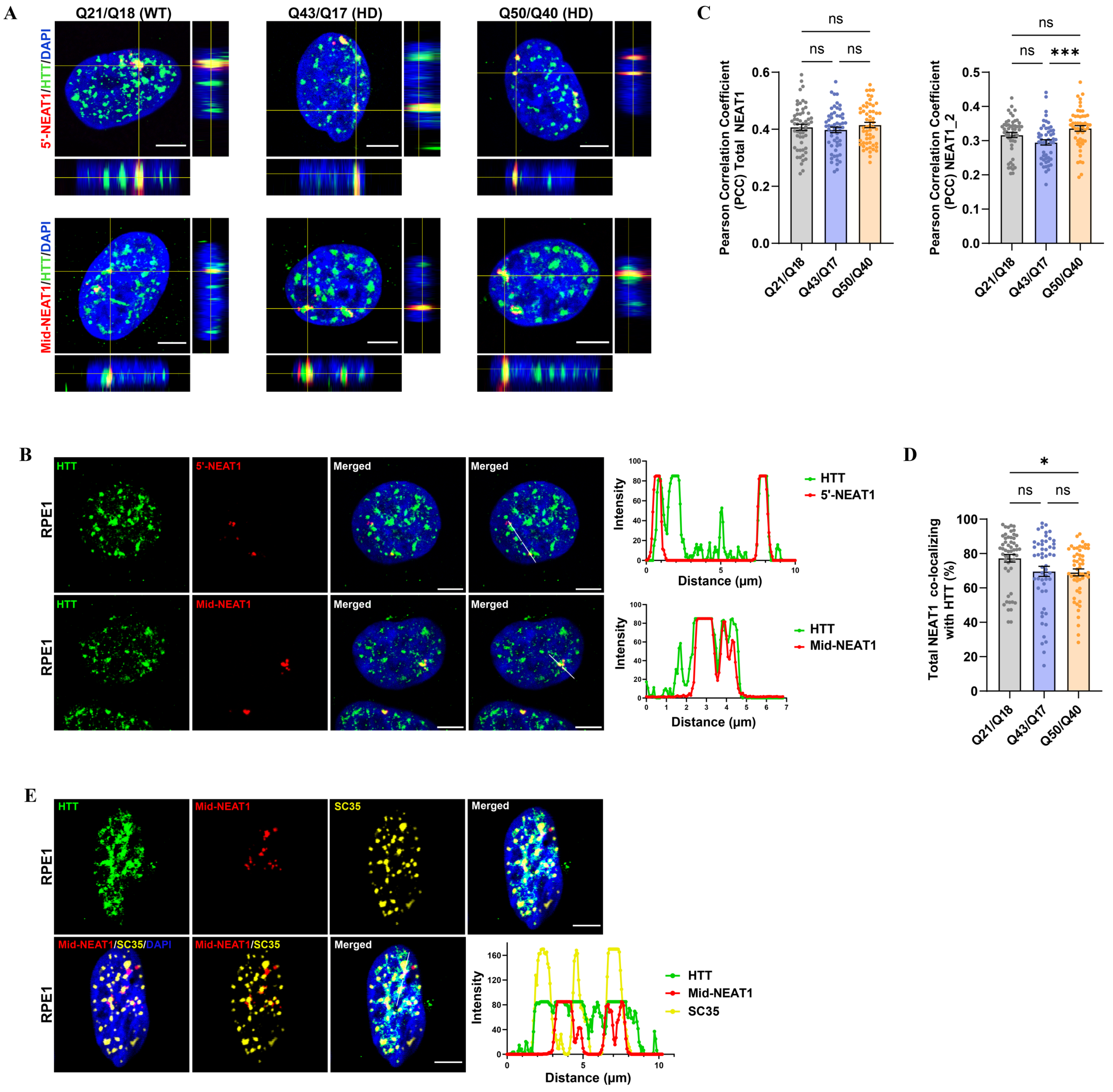


**Fig. S9. NEAT1 lncRNA and HTT protein co-localization (A)** Co-localization of HTT (green) and NEAT1 (Red), for all sets of orthogonal view images for WT and HD fibroblasts, the large image shows the x–y view, the bottom image shows the z–x and the right image shows the z–y. Scale bar indicates 5µm. **(B)** Confocal images showing co-localization of HTT (green) and NEAT1 (Red) in RPE1 cells. Quantitation of HTT and NEAT1 co-localization (performed using ImageJ). Scale bar indicates 5µm. **(C)** The graph shows Pearson's correlation coefficient for HTT and NEAT1 co-localization analysis in WT and HD fibroblasts. JACoP in ImageJ were used, and images were processed using ImageJ. Data represent mean ± sem; *n*=3. **(D)** The percentage of total NEAT1 co-localized with HTT in WT and HD fibroblasts was quantified using the JACoP in ImageJ. Data represents mean ± sem. Data were analyzed by ordinary one-way ANOVA with multiple comparisons. ∗∗∗P < 0.001 and ∗P < 0.05, ns, not significant. **(E)** Co-localization of HTT (green) with NEAT1_2 (red) and SC35+ nuclear speckles (yellow) in RPE1 cells. Quantitation of

HTT, NEAT1 and SC35+ nuclear speckles co-localization (performed using ImageJ). Scale bar indicates 5μm.
